# MCM10 compensates for Myc-induced DNA replication stress in breast cancer stem-like cells

**DOI:** 10.1101/2020.07.20.211961

**Authors:** Takahiko Murayama, Yasuto Takeuchi, Kaoru Yamawaki, Toyoaki Natsume, Rojas-Chaverra N. Marcela, Tatsunori Nishimura, Yuta Kogure, Asuka Nakata, Kana Tominaga, Asako Sasahara, Masao Yano, Satoko Ishikawa, Tetsuo Ohta, Kazuhiro Ikeda, Kuniko Horie-Inoue, Satoshi Inoue, Masahide Seki, Yutaka Suzuki, Sumio Sugano, Takayuki Enomoto, Masahiko Tanabe, Kei-ichiro Tada, Masato T. Kanemaki, Koji Okamoto, Arinobu Tojo, Noriko Gotoh

**Affiliations:** Division of Molecular Therapy, Institute of Medical Science, The University of Tokyo, Minato-ku, Tokyo, Japan; Division of Cancer Cell Biology, Cancer Research Institute, Kanazawa University, Kanazawa City, Ishikawa, Japan; Division of Cancer Differentiation, National Cancer Center Research Institute, Chuo-ku, Tokyo, Japan; Department of Obstetrics and Gynecology, Graduate School of Medical and Dental Sciences, Niigata University, Niigata, Japan; Department of Chromosome Science, National Institute of Genetics, Research Organization of Information and Systems (ROIS), Mishima City, Shizuoka, Japan; Department of Computational Biology and Medical Sciences, Graduate School of Frontier Science, The University of Tokyo, Kashiwa City, Chiba, Japan; Department of Pediatrics, Faculdade de Medicina, Universidade de São Paulo, São Paulo, Brazil; Department of Breast & Endocrine Surgery, Graduate School of Medicine, The University of Tokyo, Bunkyo-ku, Tokyo, Japan; Department of Surgery, Minamimachida Hospital, Machida City, Tokyo, Japan; Department of Gastroenterological Surgery, Kanazawa University, Kanazawa City, Ishikawa, Japan; Division of Gene Regulation and Signal Transduction, Research Center for Genomic Medicine, Saitama Medical University, Hidaka City, Saitama, Japan; Department of Medical Genome Sciences, Graduate School of Frontier Sciences, The University of Tokyo, Kashiwa City, Chiba, Japan; Department of Genetics, SOKENDAI, Mishima City, Shizuoka, Japan

**Author notes:** Corresponding Author: Noriko Gotoh, MD, PhD, Kakuma-machi, Kanazawa City, Ishikawa 920-1192, Japan.

## Abstract

Cancer stem-like cells (CSCs) are responsible for the drug resistance of tumors and recurrence while they experience DNA replication stress. However, the underlying mechanisms that cause DNA replication stress in CSCs and how they compensate for this stress remain unclear. Here we provide evidence that upregulated c-Myc expression induces stronger DNA replication stress in patient-derived breast CSCs than in differentiated cancer cells. Our results suggest critical roles for mini-chromosome maintenance protein 10 (MCM10), which is a firing (activating) factor of the DNA replication origins, to compensate for the DNA replication stress. Expression levels of MCM10 are upregulated in CSCs and maintained by c-Myc. c-Myc-dependent collisions may take place between RNA transcription and DNA replication machinery in nuclei, thereby causing DNA replication stress. MCM10 may activate dormant replication origins close to the collisions to ensure replication progression. Moreover, patient-derived breast CSCs were dependent on MCM10 for their maintenance even after enrichment for CSCs that were resistant to paclitaxel, the standard chemotherapeutic agent. In addition, MCM10 depletion decreased the growth of cancer cells but not normal cells. Therefore, MCM10 is likely to robustly compensate for DNA replication stress and facilitate genome duplication in the S-phase in cancer cells, which is more pronounced in CSCs. We provide a preclinical rationale to target the c-Myc-MCM10 axis to prevent drug resistance and recurrence.

## Introduction

Breast cancer is the most frequently observed tumor type among women worldwide. Some breast cancer patients show poor prognosis due to resistance to therapy and tumor recurrence (Torre et al., 2015). Over the past few decades, studies have shown that a subset of cancer cells have the capacity to initiate tumors (Batlle & Clevers, 2017; Saygin, Matei, Majeti, Reizes, & Lathia, 2019). These tumor-initiating cells or cancer stem-like cells (CSCs) are resistant to conventional chemotherapeutic agents, resulting in recurrence. To improve the prognosis of breast cancer patients, CSC-targeted therapeutic strategies are urgently required. Clarification of the features of CSCs is important to develop CSC-targeting therapy to remove the cells and prevent recurrence. *In vitro* tumor spheroid formation in serum-free floating culture conditions was established to enrich CSCs (Ablett, Singh, & Clarke, 2012; Ponti et al., 2005). Researchers, including our group, have used this method to shed light on the features of CSCs (Beier et al., 2007; Clement, Sanchez, de Tribolet, Radovanovic, & Ruiz i Altaba, 2007; Hinohara et al., 2012; Murayama et al., 2016; Sansone et al., 2007; Tominaga et al., 2019). Although the features of CSCs have been studied extensively, the roles of DNA replication initiation factors in CSCs have not been studied carefully.

In preparation for cell division, the whole genome must be replicated during the S-phase of the cell cycle. To rapidly generate a complete copy of the entire genome, replication of the eukaryotic genome is initiated from thousands of origins (Fragkos, Ganier, Coulombe, & Mechali, 2015; Masai, Matsumoto, You, Yoshizawa-Sugata, & Oda, 2010). The inactive MCM2–7 helicases (composed of MCM family proteins MCM2, MCM3, MCM4, MCM5, MCM6, and MCM7) bind to numerous sites of origins of DNA replication on the genome to form pre-replicative complexes (pre-RCs) in late M- and G1-phases. Following the activation of S-phase cyclin-dependent kinase (S-CDK) in the S-phase, MCM2-7 in pre-RCs are activated to form the CDC45/MCM2-7/GINS (CMG) helicase, and only ~1/10 of the chromatin-bound MCM2-7 are converted into the CMG helicase in normal cells. Subsequent recruitment of firing (activating) factors including MCM10 activates the CMG helicase to form the replisome that includes DNA polymerases, followed by initiation of bidirectional DNA replication (Douglas, Ali, Costa, & Diffley, 2018; Gambus et al., 2006; Kanke, Kodama, Takahashi, Nakagawa, & Masukata, 2012; Looke, Maloney, & Bell, 2017; van Deursen, Sengupta, De Piccoli, Sanchez-Diaz, & Labib, 2012; Watase, Takisawa, & Kanemaki, 2012). MCM10 opens the MCM2–7 ring within CMG, creating a single-stranded DNA gate for passing one DNA strand when the CMG helicase engages in fork progression (Wasserman, Schauer, O’Donnell, & Liu, 2019). On the other hand, most of the origins remain dormant, and those pre-RCs are passively removed from DNA when the replisomes approach the dormant origins during replication progression.

DNA replication stress is defined as the stalling or slowing of replication progression due to interference with the normal replication process by a variety of mechanisms, including DNA strand breaks, lack of nucleotides, etc. (Techer, Koundrioukoff, Nicolas, & Debatisse, 2017; Zeman & Cimprich, 2014). Recently, a repair process that responds to DNA strand breaks has received much attention as a potential therapeutic target. Inhibitors of poly (ADP-ribose) polymerase (PARP), a repair enzyme for single strand breaks, are clinically used in breast cancer patients with *BRCA* mutations (Pettitt & Lord, 2019). However, in the majority of patients without *BRCA* mutations, PARP inhibitors are not clearly effective. In cancer cells, constitutive activation of oncogenes is a primary cause of replication stress (Gaillard, Garcia-Muse, & Aguilera, 2015; Kotsantis, Petermann, & Boulton, 2018; Petropoulos, Champeris Tsaniras, Taraviras, & Lygerou, 2019). Although it was reported that glioblastoma stem cells suffer from upregulated DNA replication stress (Carruthers et al., 2018), the level of DNA replication stress in other types of CSCs, including breast CSCs, remains unknown. When cells suffer from replication stress, checkpoint pathways are activated (Kotsantis et al., 2018) (Petropoulos et al., 2019) (Blow & Ge, 2009). Ataxia telangiectasia- and Rad 3-related protein (ATR) kinase, and subsequently checkpoint kinase 1 (Chk1), are phosphorylated and activated. Activated Chk1 slows down cell cycle progression in the S-phase and creates a time for the dormant origins to be activated for completion of DNA replication. Many proteins included in the aforementioned DNA replication initiation machinery work together to activate the dormant origins.

*c-Myc* is a typical oncogene that is frequently overexpressed in numerous cancer types. The transcription factor c-Myc is able to induce transcription of ~15% of whole genes in the genome (Meyer & Penn, 2008). Although transcription in the G1-phase is sequentially followed by DNA replication in the S-phase in normal cells, c-Myc overexpression disrupts the cooperation between the transcription and replication machinery (Macheret & Halazonetis, 2018). As a result, they collide on the DNA strands, leading to DNA replication stress in cancer cells.

To clarify the specific features of CSCs in this study, we examined breast cancer patient-derived primary samples. We compared whole transcriptomes of CSC-enriched spheroid cells and cultured cells in the regular adherent condition. We found that pathways contributing to c-Myc activation and DNA replication stress were upregulated in CSC-enriched spheroid cells. Our results suggest that c-Myc causes frequent collisions between the transcription and replication machinery in the nuclei. Theses collisions may be one of the major causes of higher levels of DNA replication stress in CSCs compared to differentiated cancer cells. Further, we showed that the expression levels of MCM10 were increased in CSCs and differentiated cancer cells, and that expression was higher in the former than the latter, compared to normal cells. MCM10 may compensate for such replication stress by activating the dormant origins. Moreover, we demonstrated that MCM10 plays critical roles in the maintenance of CSCs, even those that are resistant to paclitaxel, a commonly used chemotherapeutic agent for breast cancer. Furthermore, by analyzing patient-derived cancer cells (PDCs), we found that MCM10 is also essential for the growth of differentiated cancer cells. Thus, inhibition of MCM10 may be a novel therapeutic strategy that targets replication initiation in CSCs.

This study is the first to demonstrate that increased activity of c-Myc is a major cause of DNA replication stress in breast CSCs. Furthermore, a c-Myc-dependent firing factor, MCM10, which is required for activation of the replisome, is essential for CSCs, probably for compensating for DNA replication stress.

## Results

### CSCs can be enriched in the spheroid culture condition and show distinct features

To identify specific features of CSCs, we first cultured patient-derived breast cancer cells both in the spheroid condition, in which cells are cultured in sphere culture medium (SCM) on ultra-low attachment dishes, and in the normal adherent condition (Fig. 1a). Spheroid cells retain their stem cell features, such as high tumor-initiating ability and expression of stemness marker proteins (Ablett et al., 2012; Dontu, Al-Hajj, Abdallah, Clarke, & Wicha, 2003; Murayama et al., 2016). Indeed, patient-derived spheroid cells showed significantly higher tumor-initiating ability *in vivo* compared to their counterparts in the adherent condition (Fig. 1b and Supplementary Fig. 1a). In addition, western blotting revealed that the expression levels of the stemness marker proteins Nanog and Oct-4 were higher in spheroid cells than in adherent cells (Fig. 1c). Immunocytochemistry (ICC) showed that strong Nanog staining was observed in the nucleus and that more spheroid cells were strongly positive for Nanog (Fig. 1d). A subpopulation of breast CSCs is enriched in the CD24^low/−^/CD44^high^ cell population (Al-Hajj, Wicha, Benito-Hernandez, Morrison, & Clarke, 2003), and thus, we investigated this population with flow cytometry. The proportion of cells in the CD24^low/−^/CD44^high^ cell fraction was higher in spheroid cells than in adherent cells (Fig. 1e). These results indicate that breast CSCs were more abundant in spheroid cells, whereas differentiated cancer cells were more abundant in adherent cells.

**Figure 1.**
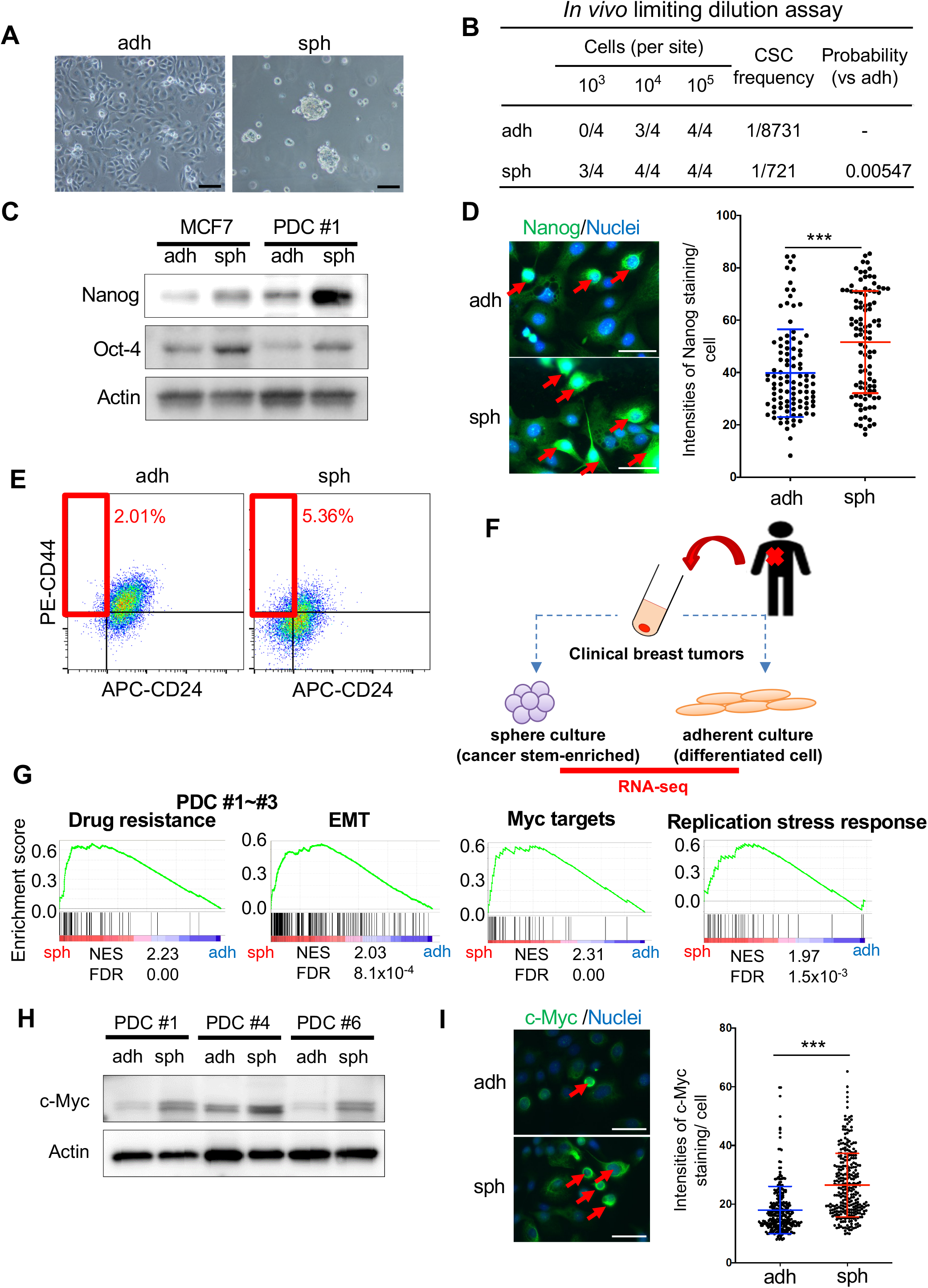
CSCs are enriched in sphere culture population and show activation of distinct pathways and c-Myc expression is upregulated in CSC-enriched spheroid cells. **a**, Images of PDC #1 cultured in adherent condition (adh; left), and in sphere culture condition (sph; right) are shown. Scale bar = 100 μm. **b**, Results of limiting dilution assay of PDC#1 obtained under adherent and sphere culture conditions were compared. CSC frequency and p-value were determined using the ELDA software (http://bioinf.wehi.edu.au/software/elda/index.html). **c**, Expression levels of MCM10 and Nanog in MCF7 and PDC #1 cells were compared between cells cultured under adherent and sphere conditions. Actin was used for loading control. **d**, (Left) Immunofluorescence images of Nanog staining in PDC #1 cells cultured under adherent and sphere conditions are shown. Nuclei were counterstained with DAPI. Arrows indicate cells with strong Nanog staining. Scale bar = 50 μm. (Right) The intensities of Nanog staining were quantified by using ImageJ software. One hundred cells in each slide were counted (mean ±SD, n = 3; ***p < 0.001). **e**, PDC #1 cells obtained by adherent and sphere culture conditions were stained with CD44 and CD24 antibodies, and then subjected to flow cytometry analysis. **f**, Schematic of the experimental procedure. Cancer cells were separated from clinical breast tumor samples, and then they were cultured in adherent and sphere conditions. RNAs were collected from both two conditions for RNA-seq transcriptome analysis. **g**, Gene set enrichment analysis (GSEA) was used to compare gene expression profiles of PDC #1-#3. Gene sets related to Drug resistance (KANG DOXORUBICIN RESISTANCE UP), EMT (SARRIO EPITHELIAL MESENCHYMAL TRANSITION UP), Myc targets (YU MYC TARGETS UP) and Replication stress response (REACTOME ACTIVATION OF ATR IN RESPONSE TO REPLICATION STRESS) were upregulated in the sphere population. NES; normalized enrichment score, FDR; false discovery ratio. **(h)** Expression level of c-Myc in PDC #1, #4 and #6 as determined by immunoblotting, were compared between cells cultured in the adherent and sphere conditions. Actin was used for loading control. **(i)** (Left) Immunofluorescence images of c-Myc staining in PDC #1 cells cultured in the adherent and sphere conditions are shown. Nuclei were counterstained with DAPI. Arrows indicate cells with strong c-Myc staining. Scale bar = 50 μm. (Right) The intensities of c-Myc staining were quantified by using ImageJ software. One hundred cells in each slide were counted (mean ±SD, n = 3; ***p < 0.001). **c, h**, Immunoblotting experiments were independently performed 3 times and representative results were presented.

Next, we performed RNA sequencing (RNA-seq) to compare the transcriptomes of spheroid cells and adherent cells (Fig. 1f). All samples were derived from the breast tumor tissues of three individual patients (PDCs #1, #2, and #3; clinical sample information is summarized in Supplementary Table 1). Gene set enrichment analysis (http://software.broadinstitute.org/gsea/index.jsp) based on the RNA-seq data revealed that genes associated with drug resistance and the epithelial-mesenchymal transition were upregulated in spheroid cells compared to adherent cells (Fig. 1g). Because these are well-known features of CSCs, we confirmed that CSCs were enriched in spheroid cells (Zhang & Weinberg, 2018). We also noticed that other gene sets related to Myc targets and the DNA replication stress response were highly upregulated in spheroid cells (Fig. 1g).

### c-Myc expression is increased in CSC-enriched spheroid cells, which may lead to strong DNA replication stress

We then investigated c-Myc protein levels. Western blotting revealed that expression levels of c-Myc were higher in spheroid cells than in adherent cells among all three PDC samples irrespective of the different subtypes (Fig. 1h, triple negative [#1], HER2 [#4], and luminal-like [#6]). ICC showed strong accumulation of c-Myc in the nucleus, and we observed that more spheroid cells were strongly positive for c-Myc than adherent cells (Fig. 1i), suggesting that a subpopulation of CSCs express c-Myc strongly.

Because gene sets related to the DNA replication stress response were upregulated in spheroid cells, we next examined the proteins involved in the checkpoint pathways (Kotsantis et al., 2018; Petropoulos et al., 2019). We found that the phosphorylation levels and amounts of ATR and Chk1 were higher in spheroid cells than in adherent cells (Fig. 2a). These results suggest that spheroid cells experienced more DNA replication stress that activated the checkpoint pathways. To directly monitor DNA replication fork stalling caused by DNA replication stress, we performed DNA fiber assays (Schwab & Niedzwiedz, 2011). We labeled cells with IdU for 30 min and then with CIdU for the next 30 min (Fig. 2b). Using this approach, bidirectional forks can be observed when a single replication origin is activated in the first 30 min (IdU: green) and then proceeds in two opposite directions (Fig. 2c, left and middle panels). If the two forks proceed normally, a CIdU-labeled (red) fork of the same length should be observed. On the other hand, when one fork stalls because of DNA replication stress, the two forks will have different lengths. We observed fork stalling more frequently in spheroid cells as reflected in the higher frequency of asymmetric CIdU-containing forks (Fig. 2c, right panel). When forks stall during the first 30 min, the stalled forks will be labeled only by IdU (green) (Fig. 2d, left panel). We observed these stalled forks more frequently in spheroid cells (Fig. 2d, right panel). Taken together, we concluded that DNA replication stress is upregulated in CSCs compared to differentiated cancer cells.

**Figure 2.**
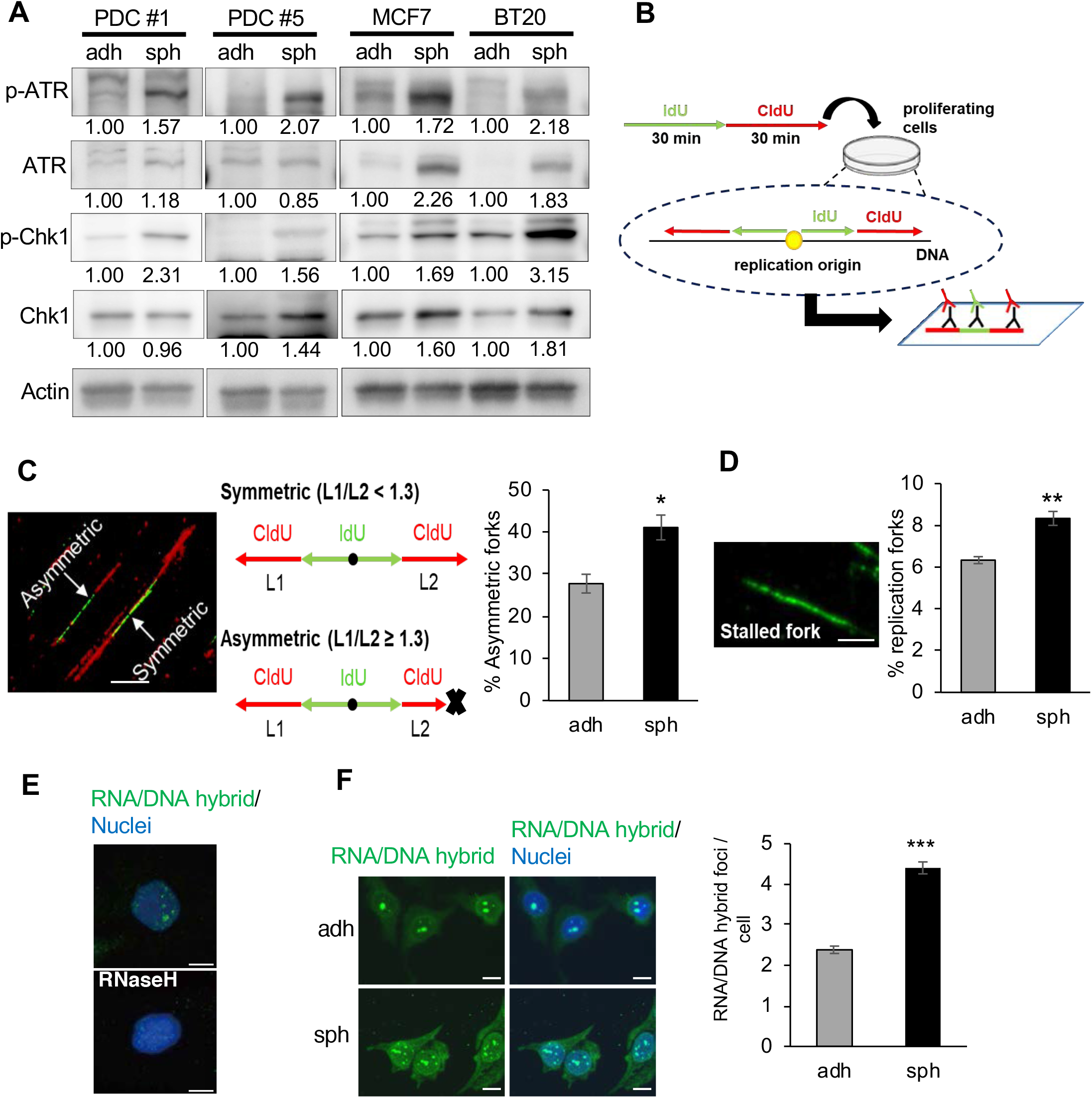
c-Myc expression and DNA replication stress are upregulated in CSC-enriched spheroid cells. **a**, Expression levels of ATR, p-ATR, Chk1 and p-Chk1 as determined by immunoblotting, were compared between cells cultured in the adherent and sphere conditions. Expression was quantified by ImageJ and normalized to Actin. The exposure time was adjusted so that the intensities of the bands were within the linear range. Experiments were independently performed 3 times and representative results were presented. **b**, Schematic of the experimental procedure. Cells were incubated sequentially with IdU then CldU. Labeled DNA was spread on glass slides, and then stained with antibodies for IdU (green) and CldU (red). If replication started in the first 30 min, bidirectional forks stained with green and red could be observed. **c**, Proportion of asymmetric forks, representative of replication stress. The ratio of longer CldU tracks (L1) to shorter tracks (L2) was calculated; forks with L1/L2 ≥ 1.3 were regarded as asymmetric. Thirty bidirectional forks in each slide were counted. Three slides for each population were prepared (mean ± SEM, n = 3; *p < 0.05). Scale bar = 10 μm. **d**, Proportion of stalled forks, labeled only with green was calculated. Two hundred labeled forks in each slide were counted. Three slides for each population were prepared (mean ± SEM, n = 3; **p < 0.01). Scale bar = 5 μm. **e**, Immunofluorescence images of RNA/DNA hybrid staining in PDCs were shown. Cells were cultured in the adherent condition with or without RNaseH treatment. Nuclei were counterstained with DAPI. Scale bar = 10 μm. **f**, (Left) Immunofluorescence images of RNA/DNA hybrid staining in PDCs cultured in the adherent and sphere conditions are shown. (Right) Number of RNA/DNA hybrid foci in each cell was counted and compared between the two conditions. Hundred cells in each slide were counted. Three slides for each population were prepared (mean ± SEM, n = 3; ***p < 0.001).

Upregulation of c-Myc induces collisions between transcription and replication machinery in the nucleus, leading to DNA replication stress (Macheret & Halazonetis, 2018). We hypothesized that upregulated c-Myc in CSCs causes collisions between transcription and replication machinery more frequently than in differentiated cancer cells. The collisions are associated with stabilized R-loops, that is, an RNA/DNA hybrid and the displaced single-stranded DNA behind elongation of RNA polymerases (Gan et al., 2011; Techer et al., 2017). R-loops can be detected with ICC staining with the monoclonal antibody S9.6, which is a widely used tool to recognize RNA/DNA hybrids (Vijayraghavan, Tsai, & Schwacha, 2016). RNA/DNA hybrid foci detected with the S9.6 antibody were localized in the nucleus (Fig. 2e). The staining disappeared following treatment with ribonuclease H (RNaseH), which cleaves RNA strands in the RNA/DNA hybrids, confirming the specificity of the antibody (Fig. 2e). We found that spheroid cells showed a higher number of RNA/DNA foci than adherent cells (Fig. 2f). These results suggest that the collisions between transcription and replication machinery occur more frequently in CSCs than in differentiated cancer cells.

We then depleted c-Myc expression by using small interfering RNAs (siRNAs) for *c-Myc* (Fig. 3a). The depletion of c-Myc led to a decreased number of RNA/DNA hybrid foci and decreased phosphorylation levels of ATR and ChK1 (Fig. 3b, c). Thus, c-Myc-induced collisions between transcription and replication machinery in the nuclei likely lead to DNA replication stress, which is more frequent in CSCs than in differentiated cancer cells.

**Figure 3.**
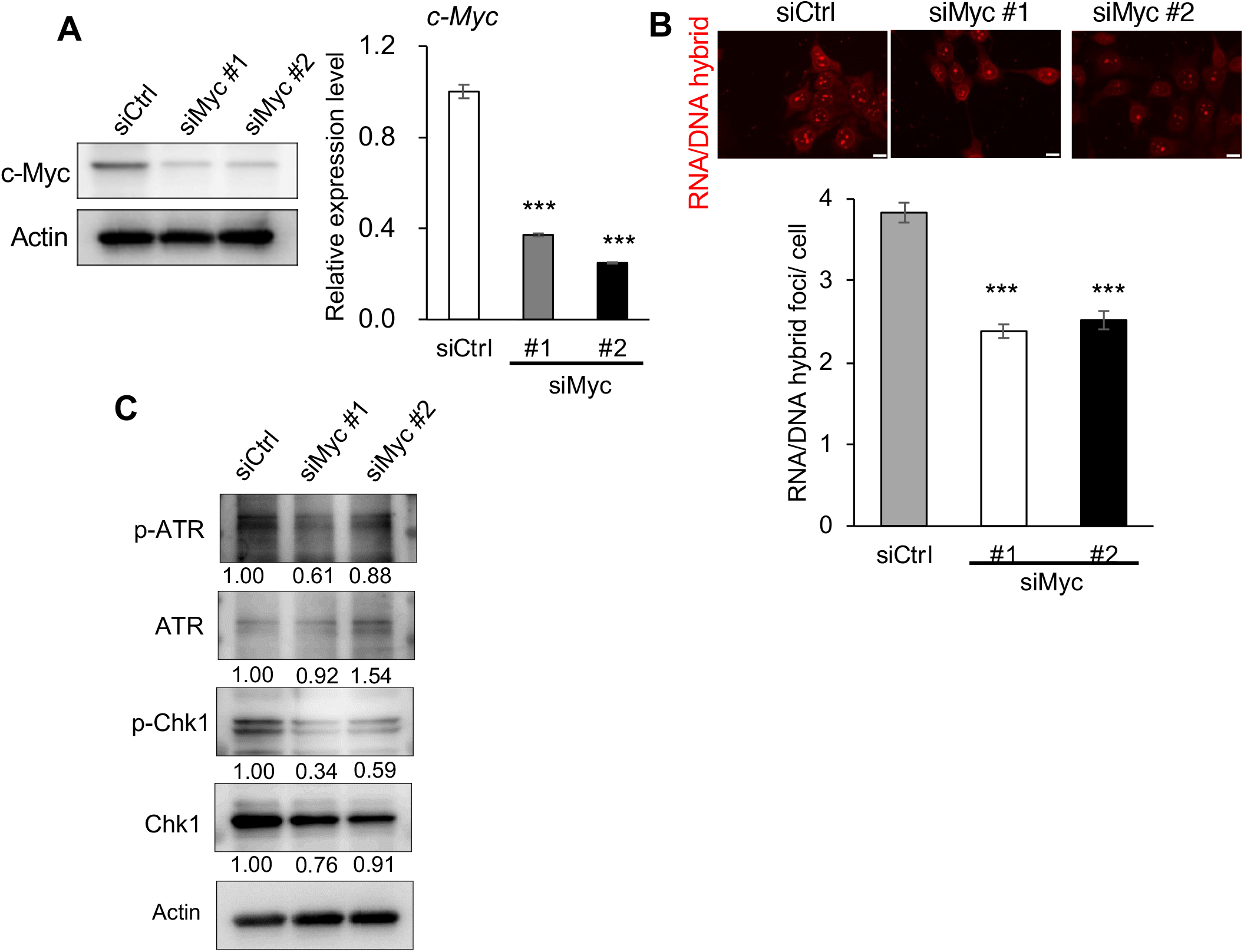
c-Myc expression contributes to replication stress. **a**, Knockdown efficiencies of siRNAs targeting *c-Myc* (siMyc #1 and #2) or control siRNA (siCtrl) in PDCs was compared by immunoblotting (left) and qPCR (right) (mean ± SEM, n = 3; ***p < 0.001). **b**, Number of RNA/DNA hybrid foci in each cell was counted and compared among spheroid cells treated with siCtrl, *siMyc* #1, and *siMyc* #2. Scale bar = 10 μm. Hundred cells in each slide were counted. Three slides for each population were prepared (mean ± SEM, n = 3; ***p < 0.001). **c**, Expression levels of ATR, p-ATR, Chk1 and p-Chk1 as determined by immunoblotting, were compared among cells treated with siCtrl, *siMyc* #1, and *siMyc* #2. Expression was quantified by ImageJ and normalized to Actin. **a,c**, Immunoblotting experiments were independently performed 3 times and representative results were presented.

### MCM10 expression is upregulated in CSC-enriched spheroid cells and co-localizes with the RNA/DNA hybrid foci in nuclei

Based on the results described above, we expected that CSCs would have mechanisms to manage higher levels of DNA replication stress. We focused our attention on *MCM10*, which was the fifth most highly upregulated gene (Supplementary Table 2), because MCM10 may activate dormant origins to compensate for DNA replication stress (Baxley & Bielinsky, 2017). Western blotting showed that the expression levels of MCM10 were higher in several breast cancer cell lines compared with MCF10A, a normal mammary epithelial cell line (Fig. 4a). We found that the expression level of MCM10 was reduced in *c-Myc*-depleted cancer cells, indicating that MCM10 expression is associated with c-Myc expression (Fig. 4b). qPCR and western blotting showed that expression levels of *MCM10* were higher in spheroid cells compared to adherent cells in several PDCs and breast cancer cell lines (Fig. 4c,d). In addition, expression of *MCM10* was higher in the CD24^low/−^/CD44^high^ cell population, a subpopulation of CSCs, than in the control population (Supplementary Fig.1b, c).

**Figure 4.**
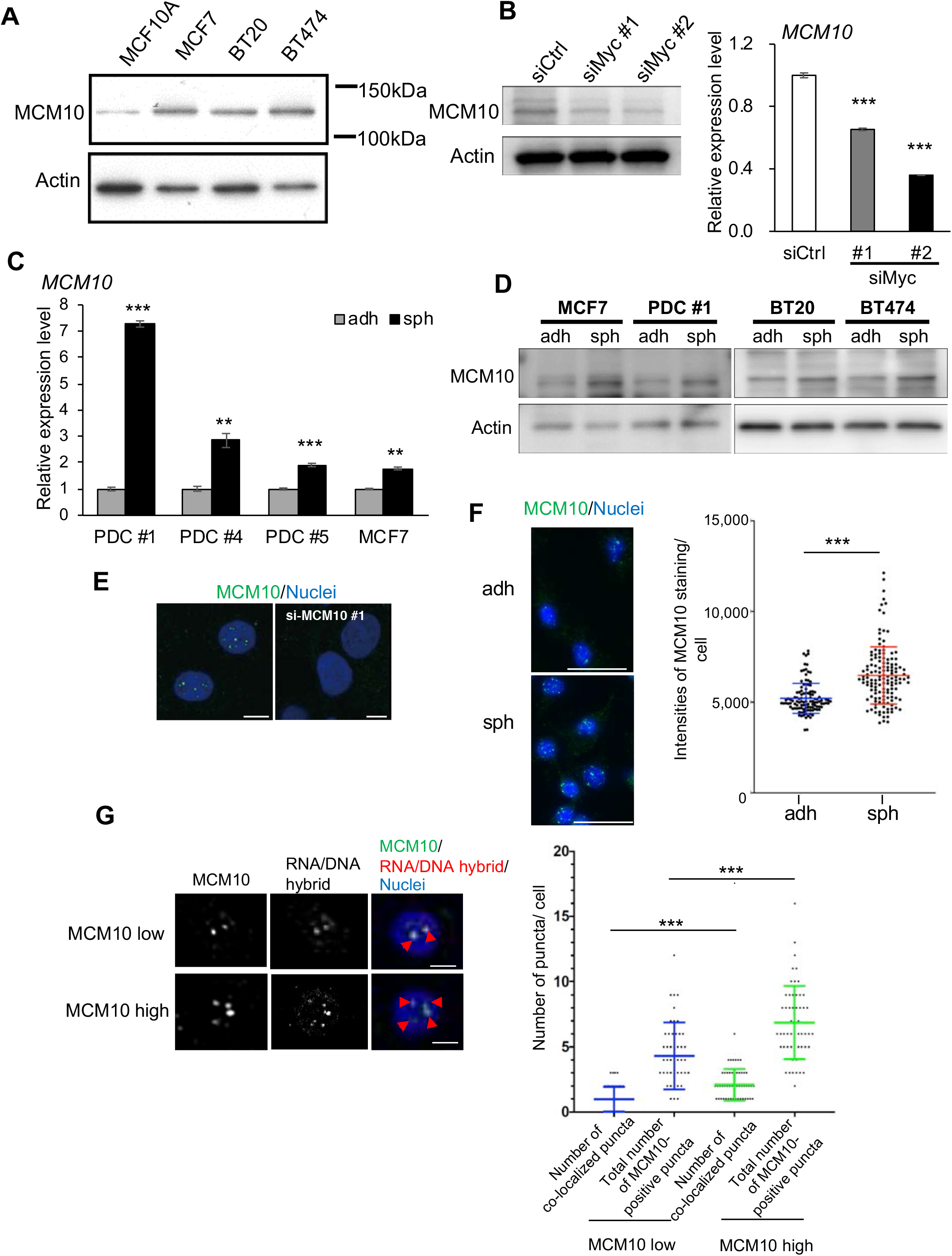
MCM10 expression is upregulated in CSC-enriched spheroid cells and MCM10 is co-localized with RNA/DNA hybrid foci. **a**, Expression levels of MCM10 in MCF10A, MCF7, BT20, and BT474 were compared by immunoblotting. Actin was used for loading control. **b**, Expression level of MCM10 in PDCs treated with siCtrl, *siMyc* #1 and *siMyc* #2 was compared by immunoblotting (left) and qPCR (right) (mean ± SEM, n = 3; ***p < 0.001). Actin was used for loading control. **c**, Expression levels of *MCM10* in PDC #1, #4, #5 and MCF7 cells were compared between cells cultured in the adherent and sphere conditions, by qPCR (mean ± SEM, n = 3; ***p < 0.001, **p < 0.01). **d**, Expression levels of MCM10 were compared in cells cultured in the adherent and sphere conditions, by immunoblotting. Actin was used for loading control. **e**, Immunofluorescence images of MCM10 staining in PDCs after transfection with control siRNA (left) or *siMCM10* #1 (right) are shown. Nuclei were counterstained with DAPI. Scale bar = 5 μm. Experiments were independently performed 3 times and representative results were presented. **f**,(left) Immunofluorescence images of MCM10 staining in PDCs cultured in the adherent and sphere conditions are shown. Nuclei were counterstained with DAPI. Scale bar = 50 μm. (right) The intensities of c-Myc staining were quantified by using ImageJ software. One hundred cells in each slide were counted (mean ±SD, n = 3; ***p < 0.001). **g**, (Left) Immunofluorescence images of MCM10 and S9.6 antibody staining in PDCs cultured in the sphere conditions are shown. The median values of the intensities of MCM10 staining (**f**) were used as the cut-off to determine MCM10-low cells and MCM10-high cells. Nuclei were counterstained with DAPI. Arrowheads indicate double positive puncta. Scale bar = 5 μm. (Right) Scatter plot showing total number of MCM10-positive puncta and double positive puncta in each cell. MCM10-positive puncta and double positive puncta were quantified by using ImageJ software. Fifty cells were counted for each group. (mean ± SD; ***p < 0.001). **a,b,d**, Immunoblotting experiments were independently performed 3 times and representative results were presented.

Further, ICC staining showed that MCM10-positive puncta were present in the nucleus (Fig. 4e). The staining disappeared by depletion of MCM10 by using siRNAs for *MCM10*, confirming the specificity of the antibodies (Fig. 4e and Fig. 5b). The number of MCM10-positive puncta were significantly higher in spheroid cells compared to adherent cells (Fig. 4f). All these results suggest that MCM10 expression is maintained by c-Myc and is upregulated in CSCs.

**Figure 5.**
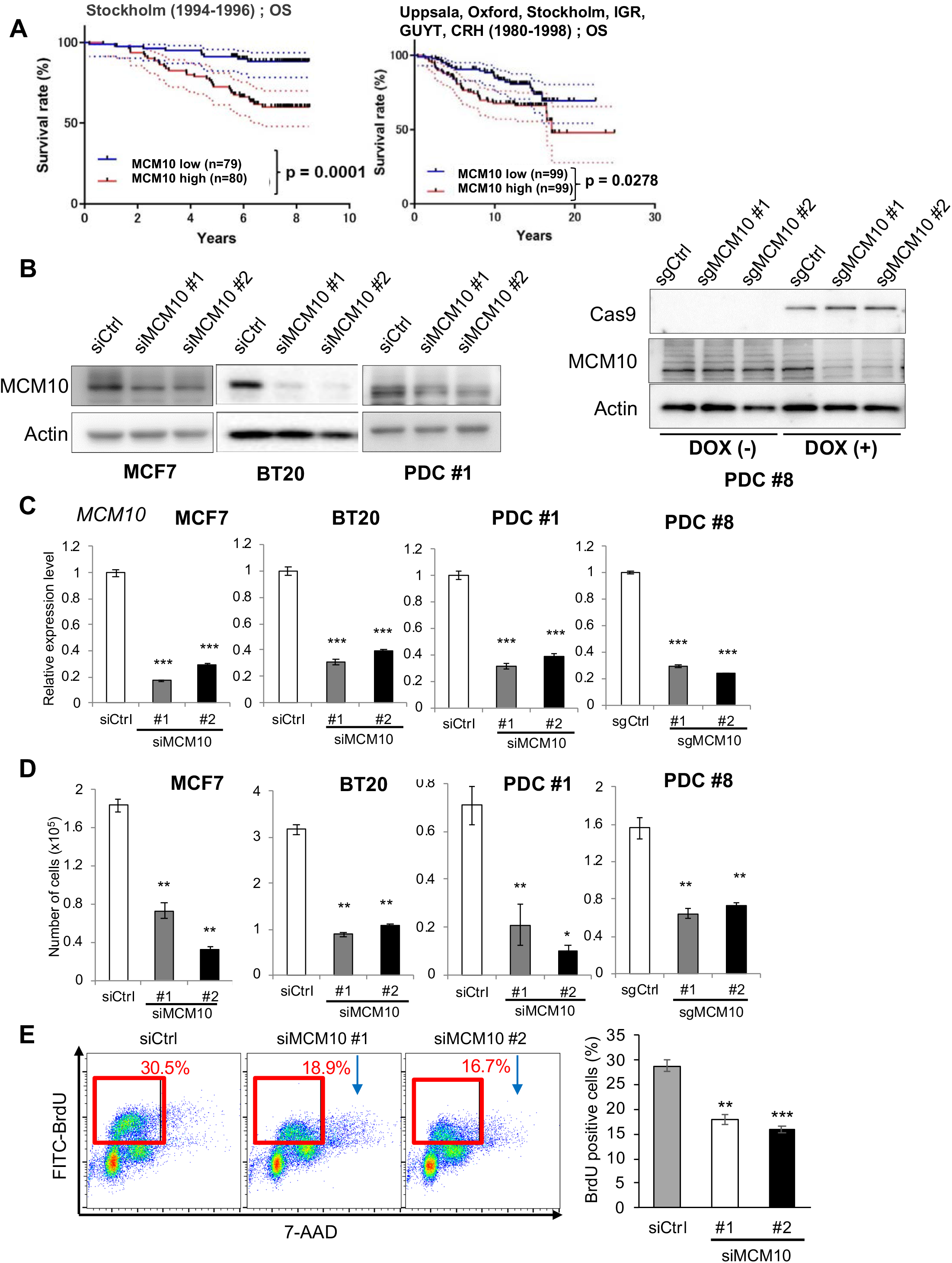
MCM10 plays important roles for proliferation of cancer cells. **a**, Kaplan–Meier survival curves were drawn using the Stockholm cohort (GSE1456; overall survival) and the Uppsala, Oxford, Stockholm, IGR, GUYT, and CRH cohorts (GSE7390; overall survival). The median values were used as the cut-off. P-values were obtained by log-rank test. **b,c**, Knockdown efficiencies of siRNAs targeting *MCM10* (siMCM10 #1 and #2) in MCF7, BT20 and PDC #1, and DOX-inducible knockout in PDC #8 were compared by immunoblotting (**b**) and qPCR (**c**) (mean ± SEM, n = 3; ***p < 0.001). Immunoblotting experiments were independently performed 3 times and representative results were presented. **d**, Cells were seeded in 12-well plates (10,000 cells/well) and cultured. Then they were harvested and counted after 4 days (mean ± SEM, n = 3; **p < 0.01, *p < 0.05). **e**, (Left) PDCs treated with siCtrl, *siMCM10* #1, or *siMCM10* #2 were incubated with BrdU for 30 min. DNA and incorporated BrdU content were analyzed by flow cytometry. (Right) Proportion of BrdU positive cells was averaged from three biological replicates (mean ± SEM, n = 3; ***p < 0.001, **p < 0.01).

To test whether MCM10 is recruited close to the collisions between the transcription and replication machinery, we stained PDCs by using the antibodies against MCM10 and S9.6. We found significant co-localization of MCM10-positive puncta and RNA/DNA hybrid foci (Fig. 4g). This result supports the notion that MCM10 is recruited to stalled forks due to the collisions in nuclei. Together, our findings suggest that MCM10 expression is upregulated in both CSCs and differentiated cancer cells, and that expression is higher in the former than the latter, compared to normal cells. Expression of MCM10 may be partly induced by upregulated c-Myc expression. Furthermore, MCM10 may be recruited to stalled forks in nuclei.

### MCM10 expression levels are prognostic, and MCM10 plays important roles in proliferation of adherent cells

To examine the clinical relevance of MCM10, we analyzed data obtained from publicly available gene expression profiles of breast cancer tissues (Desmedt et al., 2007; Pawitan et al., 2005). We found that breast cancer patients with high levels of *MCM10* expression had poor prognosis (Fig. 5a), supporting the possibility that MCM10 plays important roles in tumorigenesis. According to the Oncomine database (https://www.oncomine.org), *MCM10* expression was higher in various cancer tissues, including breast and colon cancer, than in their normal counterparts (Supplementary Fig. 2a).

To examine the functions of MCM10 in cancer cells, we depleted MCM10 with siRNAs. We confirmed that two kinds of siRNAs efficiently suppressed expression compared to the control siRNA (siCtrl) (Fig. 5b,c). We evaluated the effect of these siRNAs on proliferation of adherent cells. Knockdown of *MCM10* greatly decreased proliferation of MCF7 (luminal), BT20 (triple negative), and PDC #1 (triple negative) cells relative to a nonspecific control siRNA (siCtrl) (Fig. 5d), indicating that MCM10 is essential for the proliferation of differentiated cancer cells regardless of breast cancer subtype. To verify the requirement of high expression of MCM10 for proliferation in ovarian cancer cells, another type of gynecological cancer cell, we utilized the CRISPR-caspase 9 (Cas9)-mediated conditional knockout system to deplete *MCM10* in patient-derived ovarian cancer cells (PDC#8) (Fig. 5b right panels and c). We found that doxycycline (Dox)-induced depletion of *MCM10* led to a great reduction in cell proliferation of the ovarian cancer cells (Fig.5d). In contrast, knockdown of MCM10 in normal MCF10A cells did not significantly alter proliferation (Supplementary Fig.2b,c). These results suggest that MCM10 plays important roles in the proliferation of differentiated cancer cells but not normal cells.

We next measured DNA replication activity by examining BrdU incorporation. When *MCM10* was depleted in adherent cells, BrdU incorporation was decreased, indicating that DNA replication activity was significantly decreased in *MCM10-*depleted cells (Fig. 5e). Together, these results are consistent with the notion that MCM10 depletion increases DNA replication stress, slows down S-phase progression, and decreases cell proliferation. It appears that MCM10 is a limiting factor for dealing with DNA replication stress to complete S-phase, leading to cell proliferation.

We next examined whether MCM5, a component in pre-RCs, is a limiting factor for proliferation of cancer cells, to the same extent as MCM10. We depleted MCM5 in MCF7 cells with siRNAs. However, we found that cell proliferation was not significantly altered (Supplementary Fig.3a,b). This result indicates that MCM5 is not a limiting factor for proliferation of cancer cells in this condition. Consistently, a previous report showing that MCM2–7 is highly abundant and that MCM5-depleted cells do not show a significant growth defect in normal culture conditions (Ge, Jackson, & Blow, 2007). We then asked whether MCM10 overexpression contributes to dealing with the replication stress and thus promotes cell proliferation. To test this, we first overexpressed MCM10 and found that cell proliferation was not significantly altered by MCM10 overexpression alone (Supplementary Fig. 5c,d). We further treated cells with hydroxyurea (HU), an inhibitor of ribonucleotide reductase, to induce replication stress. Treatment with HU leads to a shortage of the deoxyribonucleotides that are used for DNA synthesis in the S-phase (Vesela, Chroma, Turi, & Mistrik, 2017). As expected, treatment with HU decreased cell proliferation, because cells experienced stronger replication stress (Supplementary Fig. 5e). We found that the decreased cell proliferation was partly restored by MCM10 overexpression when cells were treated with 500 μM HU. In this condition, we found that depletion of MCM5 blocked the restored effects on cell proliferation by MCM10 overexpression (Supplementary Fig. 5f). These results suggest that MCM10 overexpression deals with the HU-induced strong replication stress in cooperation with MCM5.

### MCM10 plays important roles in CSC properties

Next, we focused on the association between MCM10 upregulation and CSCs. To this end, we first examined the effects on tumor sphere-forming capacity. *MCM10* depletion greatly decreased the sphere-forming ability of all tested cancer cells (Fig. 6a). We subsequently examined the proportion of CD24^low/−^/CD44^high^ cells, a subpopulation of the breast CSCs. The proportion of CD24^low/−^/CD44^high^ cells was lower in *MCM10*-depleted cells than in control cells (Fig. 6b). Furthermore, the expression levels of Nanog and Oct-4 were lower in *MCM10*-depleted cells than in control cells (Fig. 6c). These results suggest that MCM10 plays an important role in the maintenance of CSCs.

**Figure 6.**
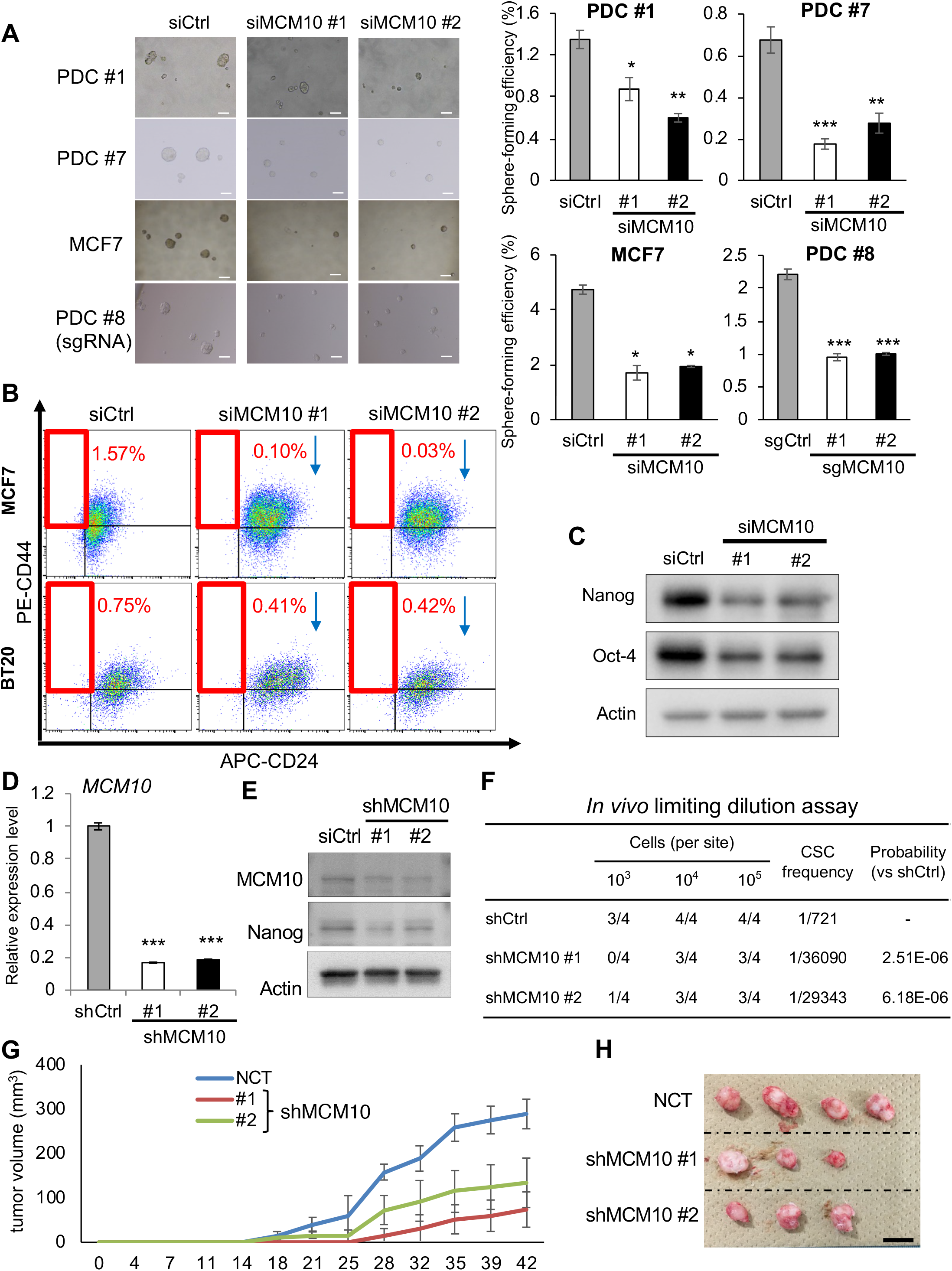
MCM10 plays important roles for CSC properties. **a**, (Left) Representative images of tumor spheres. PDC #1, #7 and MCF7 cells treated with siRNA targeting *MCM10* and DOX-inducible *MCM10* knockout PDC #8 cells were cultured under sphere conditions. Scale bar = 100 μm. (Right) Quantification of tumor sphere formation efficiency. Spheres were formed for 6 days (mean ± SEM, n = 4; ***p < 0.001, **p < 0.01, *p < 0.05). **b**, MCF7 and BT20 cells treated with siCtrl, *siMCM10* #1 or *siMCM10* #2 were stained with CD44 and CD24 antibodies, and then subjected to flow cytometry analysis. **c**, Expression levels of Nanog and Oct-4, as determined by immunoblotting, were compared between PDCs treated with siCtrl, *siMCM10* #1 and *siMCM10* #2. **d**, Expression level of *MCM10* was analyzed by qPCR in PDCs introduced with shCtrl, *shMCM10* #1, or *shMCM10* #2 (mean ± SEM, n = 3; ***p < 0.001). **e**, Expression levels of MCM10 and Nanog were compared by immunoblotting in PDCs introduced with shCtrl, *shMCM10* #1, or *shMCM10* #2. **f**, Results of limiting dilution assay of shRNA-introduced PDCs #1 were shown. Tumors larger than 50 mm^3^ were counted. CSC frequency and p-values were determined using the ELDA software. **g,h**, Growth curves (**g**) and representative images (**h**) of tumors are shown (1 × 10^5^ cells/site). Scale bar = 10 mm. **c,e**, Immunoblotting experiments were independently performed 3 times and representative results were presented.

We next constructed short hairpin RNAs (shRNAs) against *MCM10* (shMCM10 #1 and #2) and confirmed that these constructs significantly decreased the levels of MCM10 at both the RNA and protein levels relative to the control (shCtrl) (Fig. 6d, e). An *in vitro* limiting dilution assay revealed that *MCM10* depletion with shRNA decreased the tumor sphere-forming ability and estimated CSC frequency (Supplementary Fig. 3g). We then examined the tumor-initiating capacity using a patient-derived xenograft model. Using the *in vivo* limiting dilution assay, we found that *MCM10*-depleted cells had greatly reduced tumor-initiating ability and estimated CSC frequency (Fig. 6f). The tumor growth rate following inoculation of 10^5^ cells was also decreased by *MCM10* depletion (Fig. 6g, h). Together, these results illustrate the importance of MCM10 in the maintenance of CSCs *in vitro* and *in vivo*.

### Expression of c-Myc and MCM10 is enriched in paclitaxel-resistant CSCs that are dependent on MCM10 for their maintenance

Finally, we examined the possibility that *MCM10* depletion can eradicate CSCs. CSCs are enriched after paclitaxel treatment, as only differentiated cancer cells are efficiently killed by this chemotherapeutic drug (Y. Li, Atkinson, & Zhang, 2017; Samanta, Gilkes, Chaturvedi, Xiang, & Semenza, 2014). Indeed, when we treated cells with paclitaxel, the remaining resistant cells showed significantly higher sphere-forming abilities (Fig. 7a, b). In addition, we found that the expression levels of MCM10, c-Myc, and Nanog were enriched after treatment (Fig. 7c). However, when *MCM10* was depleted, the paclitaxel-resistant cells displayed a great reduction in sphere-forming abilities, and the enrichment of CSCs with high Nanog and c-Myc expression was abrogated (Fig. 7c, d). These results are consistent with the notion that CSCs that are resistant to paclitaxel can be eradicated by depletion of *MCM10*.

**Figure 7.**
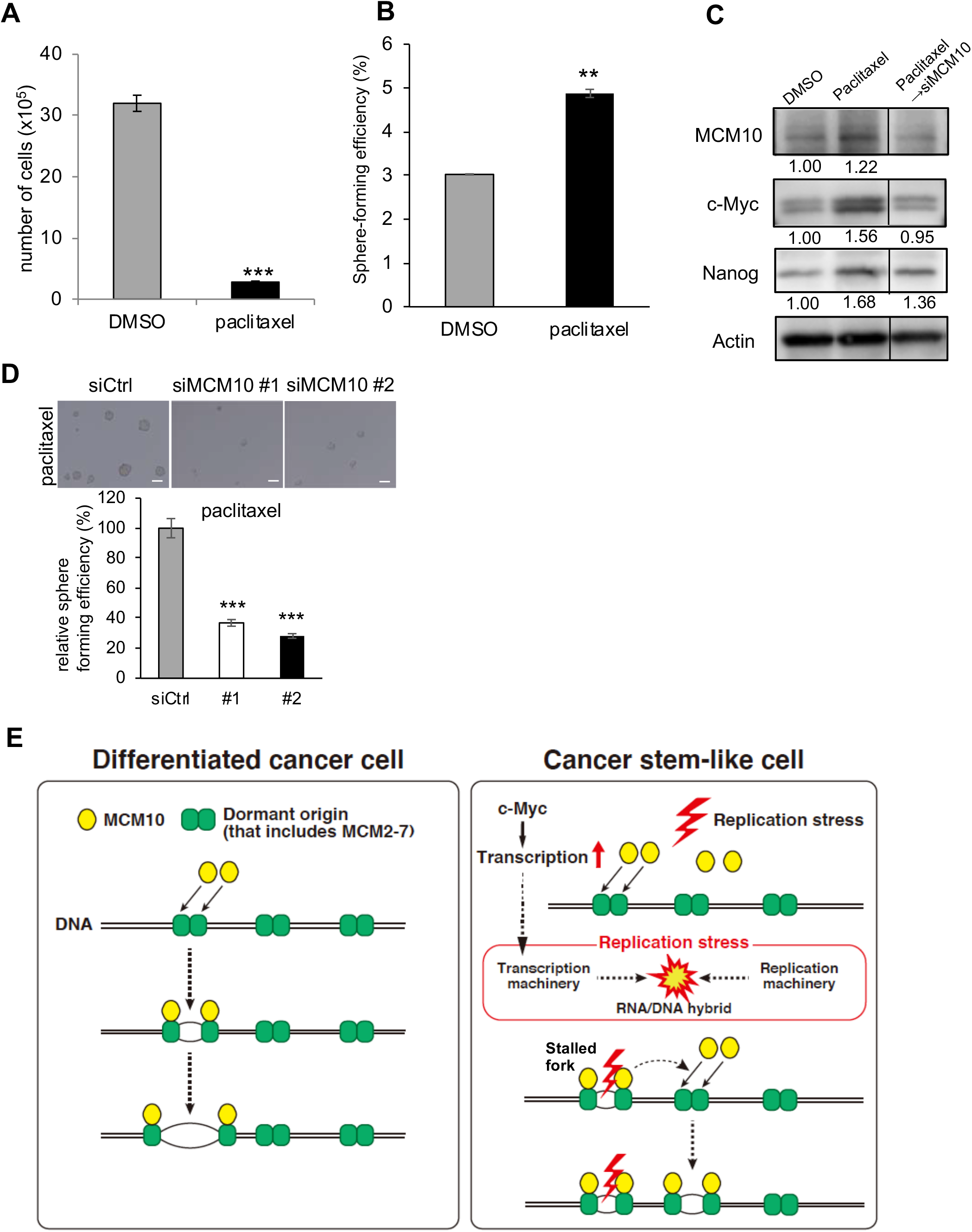
Paclitaxel-resistant cancer cells are dependent on MCM10 for their maintenance. **a**, PDCs were seeded in 12-well plates (10,000 cells/well) and cultured with 10 nM paclitaxel or control DMSO for 3 days. Cells were harvested and counted (mean ± SEM, n = 3; ***p < 0.001). **b**, Those survived cells after the treatment with paclitaxel or DMSO were used for analyzing sphere forming ability. Spheres were counted after 6 days (mean ± SEM, n = 4; **p < 0.01). **c,d**, The survived cells after the treatment with paclitaxel or DMSO were treated with siRNA for *MCM10* or control siRNA (Ctrl), and then cultured in the sphere conditions. Expression levels of Nanog, c-Myc and MCM10, as determined by immunoblotting, were compared (**c**). Expression was quantified by ImageJ and normalized to Actin. The exposure time was adjusted so that the intensities of the bands were within the linear range. Experiments were independently performed 3 times and representative results were presented. **d**, Representative images of tumor spheres are shown (upper panels). Scale bar = 100 μm. Spheres were counted after 4 days (mean ± SEM, n = 4; ***p < 0.001, **p < 0.01) (lower panel). **e**, Models of MCM10 function in CSCs and differentiated cancer cells are illustrated. In CSCs, upregulation of c-Myc leads to higher level of replication stress due to collisions between transcription machinery and replication machinery. MCM10 is necessary to deal with such DNA replication stress. MCM10 promotes completion of DNA replication by activating dormant origins near the stalled forks.

Taken together, our findings suggest that upregulated c-Myc increases RNA transcription in CSCs (Fig. 7e). The increased RNA transcription may result in collisions between transcription and replication machinery, thereby causing DNA replication stress, which is more frequent in CSCs than in differentiated cancer cells. Then, MCM10 may activate the dormant origins near the stalled replication forks. Upregulated MCM10 by c-Myc may robustly compensate for DNA replication stress by activating the dormant origins. Therefore, MCM10 is essential for the proliferation of cancer cells and maintenance of CSCs that are resistant to paclitaxel.

## Discussion

In this study, we provide evidence that a component of the DNA replication initiation machinery, MCM10, is essential for maintaining CSCs, probably by helping them compensate for DNA replication stress. MCM10 expression is upregulated in many types of cancer cells (Cui, Hu, Ning, Tan, & Tang, 2018; W. M. Li et al., 2016; Mahadevappa et al., 2018). Although previous studies reported that MCM10 upregulation is correlated with tumor malignancy, the molecular mechanisms remain largely unclear. We have provided mechanistic insight into how MCM10 is required in cancer cells, including CSCs. MCM10 is likely to efficiently activate dormant origins to compensate for DNA replication stress, which is more frequent in CSCs than in differentiated cancer cells. In contrast, the fact that depletion of MCM10 did not significantly alter cell proliferation in normal cells indicates that a small amount of MCM10 is sufficient for DNA replication in normal cells. In cancer cells, an increased amount of MCM10 appears to be required to compensate for the increased DNA replication stress. Targeting MCM10 will likely be effective for eliminating cancer cells, including CSCs, without adverse effects on normal cells.

We also show evidence that the replication stress in breast cancer cells and CSCs may be caused by collisions between the transcription and replication machinery. In neural progenitor cells and glioblastoma CSCs, transcription of long neural genes increases the frequency of collisions between the transcription and replication machinery, leading to a higher level of DNA replication stress (Carruthers et al., 2018). On the other hand, in breast CSCs, it appears that upregulated c-Myc induces the collisions between the transcription and replication machinery. Emerging evidence indicates that CSCs are maintained by unique mechanisms that may endow them with a stress-resistant phenotype relative to differentiated cancer cells. CSCs may need to produce a higher amount of proteins than differentiated cancer cells to keep their properties, possibly explaining why CSCs upregulate c-Myc expression. A better understanding of the more detailed mechanisms of replication stress in CSCs will facilitate the development of therapeutic strategies targeting CSCs.

MCM10 was the most highly upregulated gene among the DNA replication initiation factors in spheroid cells compared to adherent cells. We found that other MCMs in the pre-RCs were also among the top 100 upregulated genes. MCM5 was the 43rd most highly upregulated gene and the second most highly upregulated MCM family gene. However, MCM5 depletion did not significantly affect cell proliferation with or without treatment with HU (Supplementary Figure 3b, f) (Woodward et al., 2006). Thus, MCM10 but not MCM5 is likely essential for cancer cell proliferation. In fact, MCM2-7 helicase complexes are expressed abundantly, and cancer cells can survive after depletion of these molecules for some time (Ge et al., 2007). MCM5 remains in the pre-RCs in the licensed or dormant origins, whereas MCM10 is recruited as a firing factor to activate the origins. This functional difference between these MCMs may explain the respective phenotypes in cancer cells after knockdown of each molecule. GINS and CDC45, other firing factors, were the 12^th^ and 19^th^ most highly upregulated genes, respectively, and their roles in CSCs remain unknown. We are interested in analyzing their roles in a future project.

We provide a proof-of-principle that molecules that inhibit functions of MCM10 will be useful for targeting not only cancer cells but also CSCs. Suramin, an anti-parasitic agent, inhibits functions of MCM10 (Paulson et al., 2019). Development of inhibitors of MCM10 without enzymatic activity using Proteolysis Targeting Chimera (PROTAC) technology may be possible (An & Fu, 2018). Furthermore, the combination of such MCM10 inhibitors with chemotherapeutic reagents that induce DNA replication stress is expected to synergistically target cancer cells.

## Materials and Methods

### Cell lines and cell culture

Breast cancer cell lines MCF7, BT20 and BTB474 were purchased from the American Type Culture Collection (ATCC). Cells were cultured in RPMI1640 (GIBCO, Waltham, MA) supplemented with 10% fetal bovine serum (FBS; GIBCO) and 1% penicillin–streptomycin (P/S; Nacalai tesque, Inc., Kyoto, Japan). HEK293T cells (ATCC) were cultured for virus production in Dulbecco’s Modified Eagle Medium: Nutrient Mixture (DMEM) (GIBCO) supplemented with 10% FBS and 1% P/S. The cells were maintained in a humidified incubator with 5% CO_2_ at 37°C. They are routinely tested for contamination of mycoplasma by using PCR Micoplasma Test Kit (Takara Bio Inc., Shiga, Japan) and confirmed to be negative before performing experiments.

### Primary Cell Culture

To isolate lineage-negative (Lin^−^) breast cancer cells, cells obtained from breast tumor specimens were incubated with a mixture of biotin-conjugated antibodies against Lin^+^ cells, as previously described (Tominaga et al., 2019). The antibody mixture included a magnetic cell separation (MACS) lineage kit for depletion of hematopoietic and erythrocyte precursor cells (CD2, CD3, CD11b, CD14, CD15, CD16, CD19, CD56, CD123, and CD235a; Miltenyi Biotec, Birgisch Gladbach, Germany), endothelial cells (CD31, eBioscience, San Diego, CA), and stromal cells (CD140b, Biolegend, San Diego, CA). After incubation, cells were separated using the MACS system (Miltenyi Biotec). Isolated Lin^−^ breast cancer cells were cultured in Human EpiCult™-B Medium Kit medium (Stem Cell Technologies, Vancouver, Canada) containing a supplement mix, freshly prepared 0.48 μg/ml hydrocortisone (Stem Cell Technologies), 2 mM L-glutamine (Nacalai Tesque), 100 units/ml penicillin (Nakarai tesque, Inc.), and 100 μg/ml streptomycin (Nakarai tesque, Inc.). Isolated single cells were cultured in a humidified atmosphere at 37°C in 5% CO_2_, and the culture medium was changed every 2 days.

Tumor spheres were cultured as follows. Single-cell suspensions were cultured in ultra-low attachment plates and cells were grown in SCM, which consists of DMEM/F-12 (GIBCO), 20 ng/mL epidermal growth factor (Millipore, Burlington, MA), 20 ng/mL basic fibroblast growth factor (PeproTech, Cranbury, NJ), B27 supplement (GIBCO), and 2 μg/mL heparin (Stem Cell Technologies), as previously described (Tominaga et al., 2019). Adherent cells attached to regular cell culture plates were cultured in RPMI1640 (GIBCO) supplemented with 10% FBS (GIBCO) and 1% P/S (Nacalai Tesque Inc.). Patient-derived ovarian cancer spheroid cells (OVN62) were established from clinical specimens by the procedures as previously reported (Ishiguro et al., 2016).

### Tumor sphere formation assay

We previously confirmed that patient-derived breast cancer cells plated at 5,000 cells/mL yield tumor spheres clonally derived from single cells (Hinohara et al., 2012). Hence, cells were plated as single-cell suspensions on ultra–low-attachment 24-well plates (1000–5000 cells/well) to obtain single cell–derived tumor spheres. The cells were grown in SCM. Spheres with diameter >75 μm were counted after 4–7 days.

### Generation of ovarian cancer spheroid cells with inducible-CRISPR/Cas9 targeting *MCM10*

Inducible-Cas9 lentiviral plasmid (Edit-R Inducible lentiviral Cas9; CAS11229) were purchased from Horizon Discovery (Cambridge, UK). For production of lentivirus encoding inducible-Cas9 nuclease, the lentiviral plasmids and packaging plasmids were transfected into lentiX-293T cells using Lipofectamine 2000 (Invitrogen, Carlsbud, CA) and lentivirus-containing supernatants were harvested after 3 days. The lentivirus-containing media was transferred onto OVN62 cells (Ovarian cancer spheroid cells) to generate cells expressing inducible-Cas9 and incubated with blasticidin (5 μ g/mL) (Nakarai tesque, Inc.) for 3 days. After selection, OVN62 cells heterogeneously expressing inducible-Cas9 were performed single-cell sorting using a FACS Aria Ⅲ Cell Sorter (BD Bioscience, San Jose, CA) to pick up stably expressing inducible-Cas9 construct.

We performed a modification of pLenti-sgRNA plasmid (Addgene #71409) with the sgRNA scaffold with the sgRNA sequence targeting *MCM10* (MCM10 sgRNA #1: 5’-CGGTGAATCTTATACAGAAG-3’, MCM10 sgRNA #2: 5’-GAGGGTGGCTCGAACACCAA-3’, MCM10 sgRNA #3: 5’-CGGTGAATCTTATACAGAAG-3’ and 5’-GAGGGTGGCTCGAACACCAA-3’). OVN62 with stably expressing inducible-Cas9 and targeting *MCM10* were selected by puromycin selection (2 μg/mL) (Nacalai tesque, Inc.).

### Cell viability assay for ovarian cancer spheroid cells

OVN62 cells stably expressing inducible-Cas9 nuclease and *MCM10*-targeting sgRNA (Non-target sgRNA, MCM10 sgRNA #1, and #2) were treated with Doxycycline (Dox) (Nacalai tesque, Inc.) for 3 days to express Cas9 and induce *MCM10* knockout. After DOX treatment, cells were dissociated to single cells and seeded 3000 cells in each well of the 96-well plates. After 0, 3, 7 days incubation, cell viability was measured by using CellTiter-Glo Assay (Promega, Madison, WI).

### RNA extraction, cDNA amplification, library preparation, and sequencing

Total RNA was extracted from cells using the NucleoSpin RNA XS kit (Clontech, Moutain View, CA). The Smarter Ultra low RNA input kit (Clontech) was used for the synthesis and amplification of cDNA using up to 10 ng of total RNA following the manufacturer’s instructions and performing no more than 12 cycles of PCR in order to minimize amplification biases. The quality of cDNA was verified by Agilent 2100 Bioanalyzer using High Sensitivity DNA Chips (Agilent Technologies, Santa Clara, CA). Truseq DNA Illumina libraries were prepared and sequenced to obtain approximately 90 million reads (101 bp paired-end reads) per library using the Hiseq 2000/2500 Illumina sequencer (San Diego, CA).

### RNA-sequence data analysis

Sequences were trimmed to remove adaptors and low-quality bases. Trimmed reads were mapped onto the hg19 genome (UCSC human genome 19, version:20150519) using Tophat 2.0.10 and transcripts were assembled by Cufflinks 2.1.1 based on RefSeq gene annotation. Transcript expression levels were quantified by Cuffdiff 2.1.1 using the fragments per kilobase of transcript per million mapped fragments (FPKM) method.

GEO accession number is GSE127264.

### Real-time PCR analysis

Total RNAs were extracted using TRIzol Reagent (Thermo Fisher Scientific, Waltham, MA) according to the manufacturer’s instructions. The High-Capacity cDNA Reverse Transcription Kit (Thermo Fisher Scientific) was used to prepare the cDNA solution. For real-time PCR analyses, Taqman probes of *RPF1*, *LTV1*, *ESF1*, *NSA2*, *BRIX1*, *FCF1*, *MCM10* and *Nanog* were purchased from Applied Biosystems. For detecting premature RNA, primer sequences designed by Kofuji et al. (Kofuji et al., 2019) (pre-rRNA F; 5’-TGTCAGGCGTTCTCGTCTC-3’, pre-rRNA R; 5’-AGCACGACGTCACCACATC-3’) were used. Reactions were performed using the pre-set program of the ABI ViiA 7 Real-Time PCR System (Thermo Fisher Scientific).

### siRNA

We purchased two different siRNA duplexes of *MCM10* (#1, HSS124480 and #2, HSS124482), two different siRNA duplexes of *Myc* (#1, VHS40785 and #2, VHS40789) and a nonspecific control siRNA duplex with similar GC content (siCtrl; Medium GC Duplex #2) from Invitrogen. siRNAs against *MCM5* were designed according to a previous report (Ge et al., 2007); target sequences were 5’-GGAGGUAGCUGAUGAGGUGTT-3’ (#1) and 5’-AAGCAGUCGCAGUGAAGAUUG-3’ (#2). siRNAs were transfected using RNAiMAX (Invitrogen).

### Transient overexpression of MCM10

Cells were transfected with pCMV6-Myc-DDK-MCM10 (OriGene, Rockville, MD) and control vector using ViaFect Transfection Reagent (Promega, Madison, WI).

### Western blot analysis

Immunoblotting was performed using standard procedures as described (Hinohara et al., 2012). Antibodies against Nanog (4903S), Oct-4 (2750S), ATR (2790S), p-ATR (2853S), Chk1 (2360T), p-Chk1 (2349T), c-Myc (5605S) and Myc-tag (2278) were purchased from Cell Signaling Technology (Danvers, MA). Antibodies against MCM10 (3733) and MCM5 (17967) were purchased from Abcam (Cambridge, UK). Anti-actin antibody (MAB150) was purchased from Millipore. Cas9 antibody was purchased from Active Motif. Proteins were detected with horseradish peroxidase–conjugated anti-mouse or anti-rabbit antibodies (GE Healthcare Life Sciences, Marlborough, MA).

### Immunocytochemistry

Cells in adherent and sphere culture condition were plated on BioCoat Culture Slide (Corning, Corning, NY) after trypsinization, and incubated for 6 h. To detect expression of proteins, cells were fixed with 4% paraformaldehyde (PFA) (Wako, Osaka, Japan) or 100% methanol (Wako, Osaka, Japan), 6 h after seeding, the shortest period for cell attachment. Cells were incubated with 0.2% Triton X-100 (Wako, Osaka, Japan) to permeabilize membranes, and stained overnight with primary antibodies and for 1 h with secondary antibodies. Immunofluorescent visualization of Nuclei was counterstained with DAPI (Thermo Fisher Scientific). Coverslips were mounted with Fluorescence Mounting Medium (Dako, Glostrup, Denmark). Immunofluorescence was detected using an Olympus IXplore pro microscope (Tokyo, Japan) or Nikon confocal microscopy (A1 HD25) (Tokyo, Japan) with the ANDOR software. Acquired images were analyzed by ImageJ software. Antibodies against Nanog (4903S) and c-Myc (5605S) were purchased from Cell Signaling Technology. MCM10 antibody was purchased from Invitrogen (PA5-67218). S9.6 antibody that binds to RNA/DNA hybrid was purchased from Millipore (MABE1095).

### RNaseH treatment

Cells were incubated with Ribonuclease H RNaseH (60 U/μl) (Takara Bio Inc., Code No. 2150A) for 4 hours before immunocytochemistry assay.

### Proliferation assay for breast cancer cells

Cells were seeded in 12-well plates at low density (5000–10000 cells/well), cultured in RPMI1640 supplemented with 10% FBS and 1% P/S or DMEM with 10% FBS and 1% P/S. HU (Wako, Tokyo, Japan) was added to the medium as necessary. After 4–6 days, cells were harvested and counted.

### Flow cytometry analysis

To identify the breast CSC population, cells were stained with Alexa Fluor 647–conjugated anti– human CD24 and APC-H7 labeled anti–human CD44 antibodies (BD Pharmingen, San Jose, CA) at 4°C for 20 min. The cells were then analyzed with a FACSAria II flow cytometer (BD Bioscience, San Jose, CA). Dead cells were excluded by propidium iodide (PI; Sigma, St. Luis, MO) staining. Data were analyzed using the FlowJo software (TreeStar, San Carlos, CA).

To detect DNA-binding MCM3, collected cells were first treated with 750 μL low-salt extraction buffer (0.1% Igepal CA-630, 10 mM NaCl, 5 mM MgCl_2_, 0.1 mM PMSF, 10 mM potassium phosphate buffer [pH 7.4]) for 5 min on ice. Then, the cells were fixed by adding 250 μL 10% formalin (SIGMA). After incubation at 4°C for 1 h, the cells were washed with phosphate buffered saline (PBS)(GIBCO). Extracted cells were then incubated with anti-MCM3 antibody (Abcam) and anti–rabbit IgG secondary antibody (Alexa Fluor 488) (Molecular Probes, Eugene, OR) in flow buffer (0.1% Igepal CA-630, 6.5 mM Na_2_HPO_4_, 1.5 mM KH_2_PO_4_, 2.7 mM KCl, 137 mM NaCl, 0.5 mM EDTA [pH 7.5]). Cells were analyzed on a FACSAria II flow cytometer (BD Bioscience) after staining with Hoechst 33258 (Sigma) to detect nuclear DNA (1 μg/mL).

### DNA fiber assay

Adherent and sphere-cultured cells were pulsed-labeled with 25 μM ldU (Sigma) for 30 min, followed by 250 μM CIdU (Sigma) for 30 min. The cells were trypsinized and resuspended in 100 μL PBS (GIBCO)(10^5^–10^6^ cells/mL). Then, a 2 μL cell suspension was placed at the end of a glass slide. After air drying for 8 min, 7 μL of fiber lysis solution (50 mM EDTA, 0.5% SDS, 200 mM Tris-HCl [pH 7.5]) was pipetted on top of the cell suspension and mixed. Cell lysis proceeded for 5 min, and then the slides were tilted at 15° to allow the DNA spread down the slide. Slides were air-dried for 15 min and fixed in methanol/acetic acid (3:1). After washing with distilled water, DNA was denatured in 2.5 M HCl for 80 min. The slides were washed with PBS three times, and blocked for 1 h in 5% bovine serum albumin (BSA) (Sigma) in PBS (GIBCO). After blocking, the slides were incubated with primary antibodies (anti-CldU, Abcam ab6326; anti-IdU, BD 347580) followed by secondary antibodies (Alexa Fluor 594–conjugated anti–rat IgG and Alexa Fluor 488–conjugated anti–mouse IgG)(Molecular Probes).

### Plasmid construction

The pLKO shRNA vector was used for knockdown experiments. Target sequences for human *MCM10* were 5’-TCATCCTCAGAAGGTCTTAAT-3’ (#1) and 5’-GGACTTAACAGATGAAGAAGA-3’ (#2). Lentiviral plasmids were transduced into HET293T cells along with ViraPower Lentiviral Packaging Mix (Invitrogen) using the Lipofectamine 3000 Transfection Reagent (Invitrogen). The medium was changed after 16 h.

### Transduction of patient-derived cancer cells with lentiviral vectors

Culture supernatant from HEK293T cells containing virus particles was applied to patient-derived cancer cells. The cells were incubated at 37°C in 5% CO_2_ for 48 h, and then virus-infected cells were selected using 2.5 μg/mL puromycin for breast cancer cells and 2 μg/mL puromycin (Nakarai tesque, Inc.) for ovarian cancer spheroid cells.

### *In vivo* limiting dilution assay

Seven-week-old female immunodeficient NSG mice were anesthetized with isoflurane (Abbott, Lake Bluff, IL). Patient-derived breast cancer cells infected with lentivirus (shMCM10 #1, #2, and shCtrl), or cells cultured in adherent and sphere condition were suspended in 50 μL Matrigel (BD Biosciences) in a dilution series (10^3^, 10^4^, and 10^5^ cells). Suspended samples were then injected subcutaneously into the mammary fat pads of NSG mice. Tumor volume was measured twice a week using the following formula: *V* = *1/2*(*L* × *W*^*2*^), where *L* equals length and *W* equals width. Tumors larger than 50 mm^3^ were counted.

### Statistical analysis

All data are presented as means ± SEM or means ± SD. The unpaired Student *t*-test was used to compare differences between two samples, and values of p < 0.01–0.05 (*), p < 0.001–0.01 (**), or p < 0.001 (***) were considered significant. Tumor-initiating frequency was calculated using the ELDA Software. Kaplan-Meier survival curves were analyzed by log-rank test.

### Study approval

All human breast carcinoma specimens were obtained from The University of Tokyo Hospital, Minami-Machida Hospital and Kanazawa University Hospital. Human ovarian cancer specimens were obtained from Niigata University Medical & Dental Hospital. This study was approved by the institutional review boards of the Institute of Medical Science, The University of Tokyo; The University of Tokyo Hospital; Minami Machida Hospital; National Cancer Center; Niigata University; and Kanazawa University. Written informed consent was received from all participants before inclusion in the study.

## Acknowledgements

We thank H. Nakauchi, Y. Ishii, and A. Fujita for their help with flow cytometry. This work was supported in part by an Extramural Collaborative Research Grant from the Cancer Research Institute, Kanazawa University, a Research Grant from Princess Takamatsu Cancer Research Fund (17-2924), a Grant-in-Aid for Scientific Research from JSPS (17K19587 and 18H02679), and a research grant from AMED Project for Development of Innovative Research on Cancer Therapeutics, Project for Cancer Research and Therapeutic Evolution (16cm0106120h0001) and Practical Research for Innovative Cancer Control (16ck0106194h0001) to N.G. This work was also supported by MEXT KAKENHI (No. 221S0002) to N.G. This work was supported by the Graduate Program for Leaders in Life Innovation (GPLLI) at The University of Tokyo Life Innovation Leading Graduate School (to T.M., http://square.umin.ac.jp/gplli/en/index.html) and a Grant-in-Aid for a Japan Society for Promotion of Science (JSPS) fellowship (to T.M., http://www.jsps.go.jp/english/e-pd/index.html).

## Author contributions

T.M. performed experiments, analyzed and interpreted data and wrote the manuscript. Y.T., K.Y., NM.RC., T. Nishimura, Y.K., A.N., K.T. and A.S. performed experiments and analyzed data. T. Natsume and M.T.K. helped fiber assay, analyzed data and provided scientific insight. M.T.K. also wrote the manuscript. M.Y., S.I, M.I., T.O., T.E., M.T. and K.T. provided clinical samples. K.I., K.H-I., S.I. and K.O. analyzed data and provided scientific insight. M.S. and Y.S. performed RNA sequencing and analyzed data. S.S. and A.T. provided scientific insight and supervised the study. N.G. conceived the study, analyzed and interpreted data and wrote the manuscript. All authors read the manuscript and provided feedback on the manuscript.

---

**Supplementary Figure 1.**
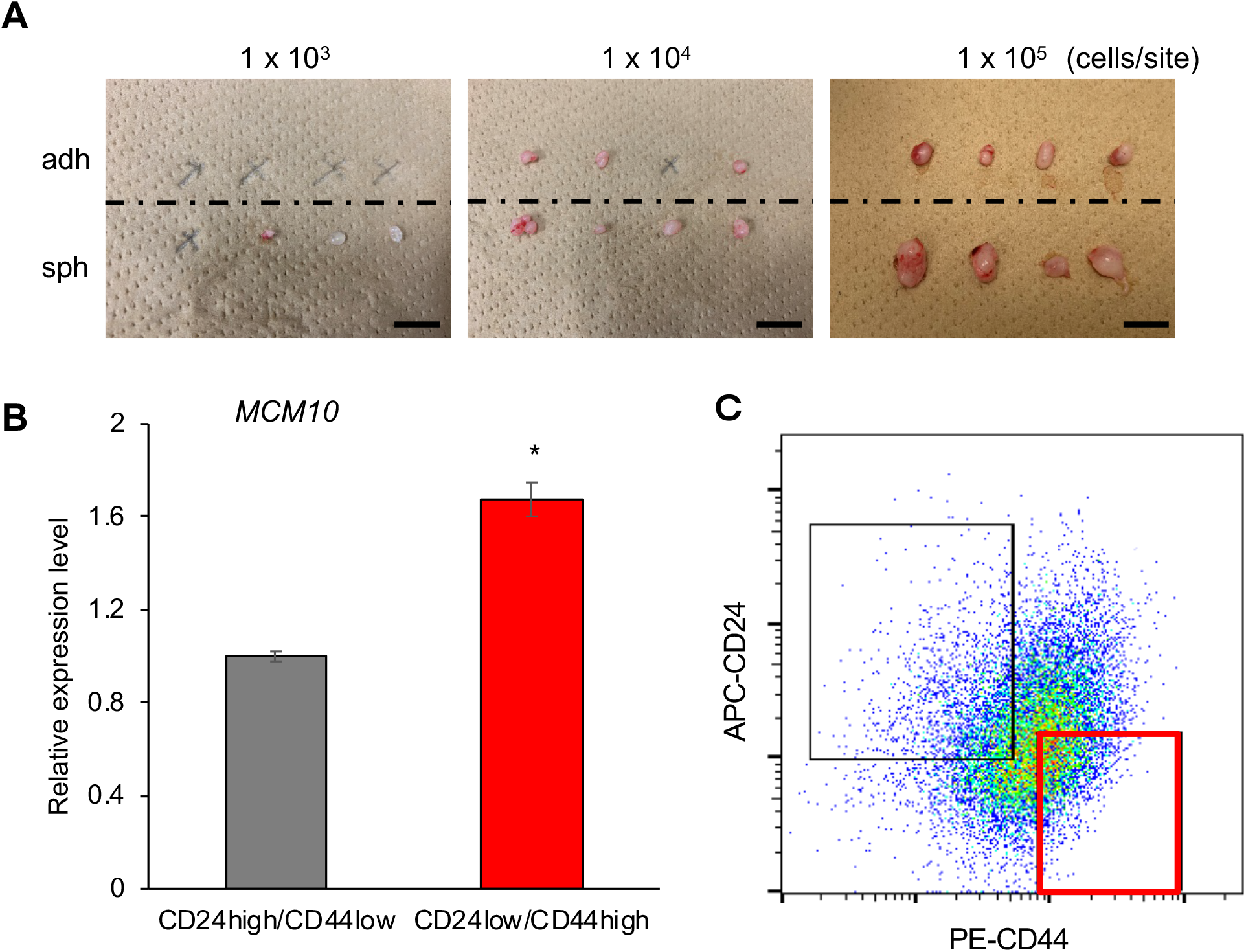
**a**, Representative images of tumors formed in the *in vivo* limiting dilution assay (1 × 103, 104 and 105 cells/site). Scale bar = 10 mm. **b**,**c**, Expression level of *MCM10* was compared by qPCR between the CD24^−/low^/CD44^high^ CSC-enriched population and the CD24^high^/CD44^low^ control population. The PDC #6 cells were analyzed. (mean ± SEM, n = 3; *p < 0.05).

**Supplementary Figure 2.**
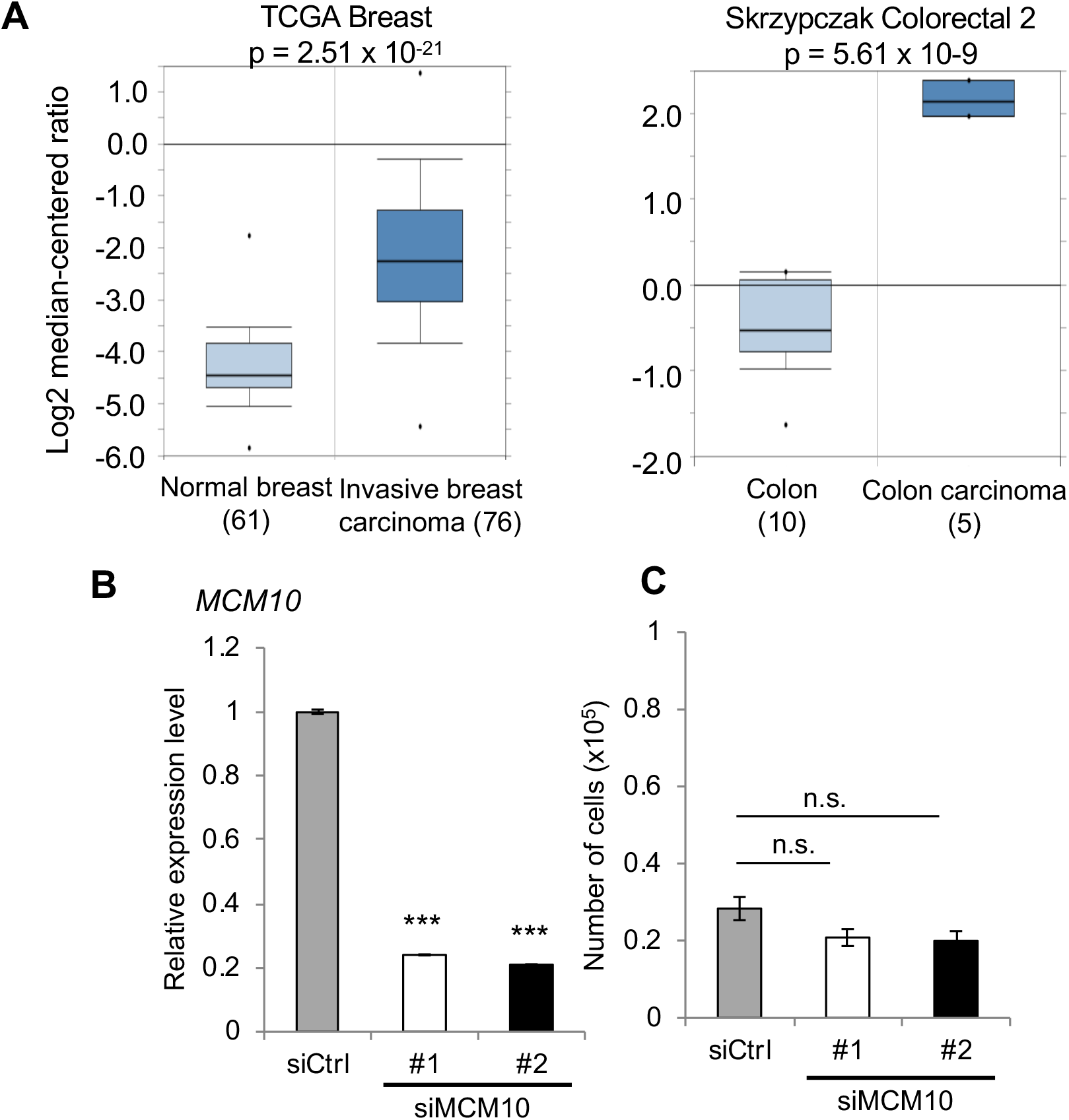
**a**, *MCM10* expression was compared between non-malignant cells and cancer cells in breast and colon using the Oncomine cancer gene expression database (Right; TCGA Breast, Left; Skrzypczak Colorectal 2). P-values were calculated by Student’s t-test. **b,c**, MCF10A treated with siCtrl or *siMCM10*. Knockdown efficiencies of siRNAs (**b**) and growth rates (**c**) were compared.

**Supplementary Figure 3.**
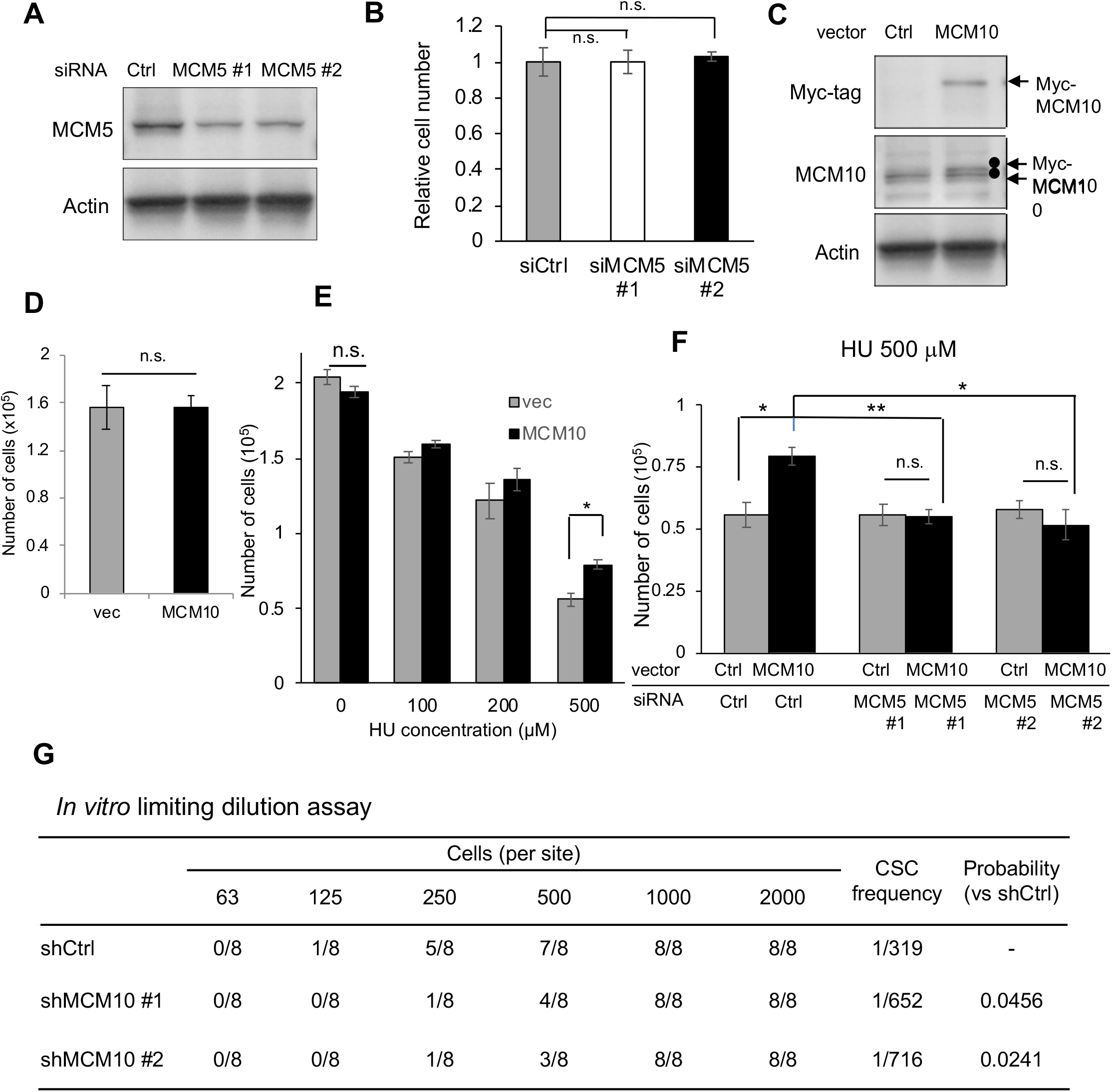
**a, b**, Expression levels of MCM5 were compared in MCF7 cells treated with siCtrl or *siMCM5* (**a**). Number of cells were counted after 4 days (mean + SEM, n = 3) (**b**). **c,d**, Expression levels of endogenous MCM10 and Myc-tagged MCM10, as determined by immunoblotting, were compared among cells transfected with the indicated expression vectors (**c**). Number of cells were counted after 4 days (mean + SEM, n = 3)(**d**). **e,f**, MCF7 cells transfected with the indicated vectors (**e**) and siRNAs (**f**) were seeded in a 12-well plate (10,000 cells/well). Forty-eight hours later, they were treated with indicated concentrations of HU for an additional 48 h. Cells were harvested and counted (mean + SEM, n = 3; **p < 0.01, *p < 0.05). **g**, In vitro limiting dilution assay for MCM10-depleted cells in patient-derived breast cancer cells: 2,000, 1,000, 500, 250, 125, or 63 cells were seeded in each well of a 96-well ultra–low-attachment plates. Results were obtained 7 days after seeding. CSC frequency and p-values were determined using the ELDA software.

**Supplementary Table 1.**
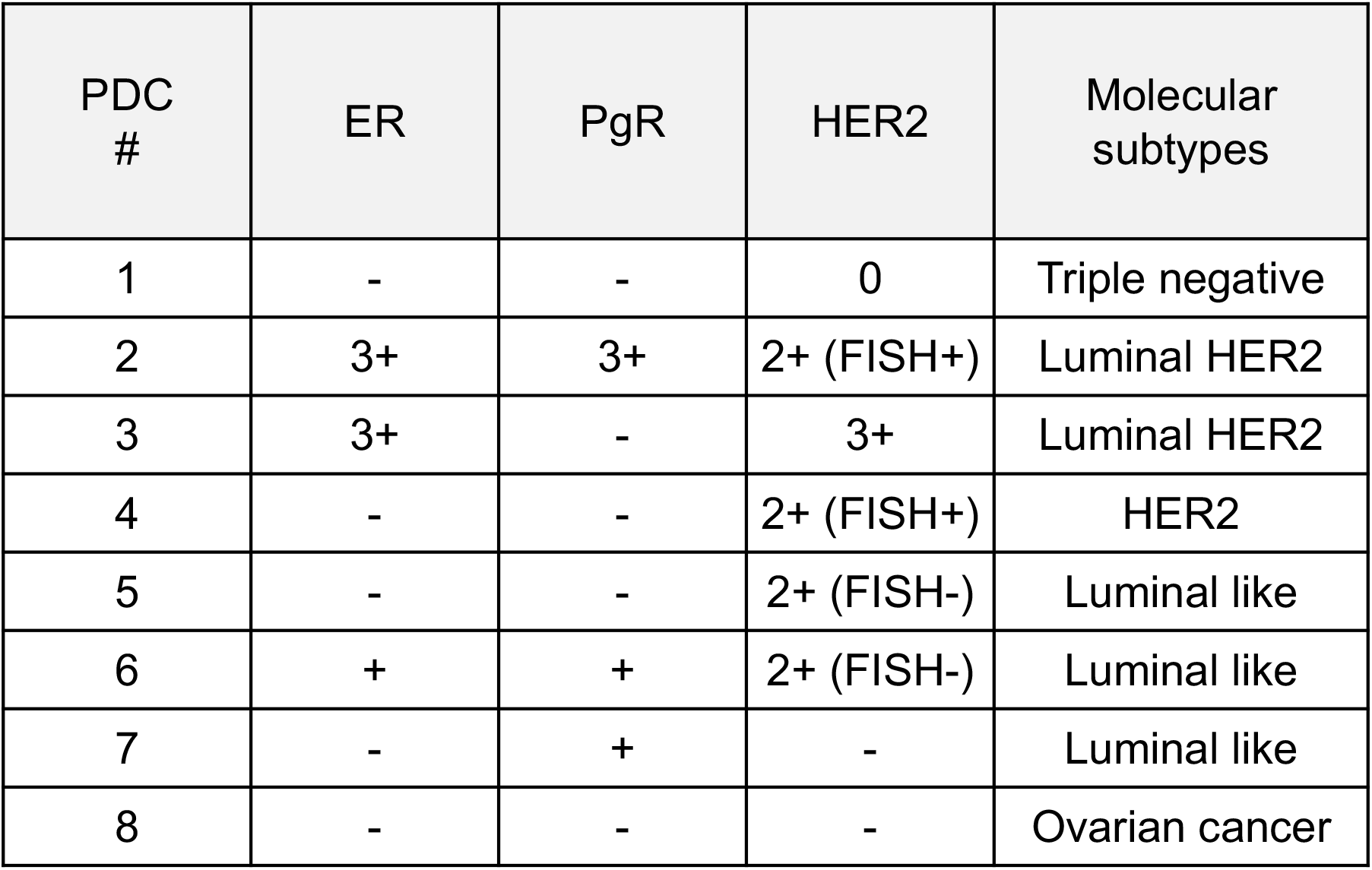
Characteristics of clinical breast tumors used in this study

**Supplementary Table 2.** Genes included in Reactome_DNA_Replication gene set and the ratios of expression levels of each gene, sphere cells (SPH) / adherent cells (ADH)

| Rank | Gene symbol | SPH/ADH | Rank | Gene symbol | SPH/ADH | Rank | Gene symbol | SPH/ADH | Rank | Gene symbol | SPH/ADH |
| --- | --- | --- | --- | --- | --- | --- | --- | --- | --- | --- | --- |
| 1 | GMNN | 37.044 | 51 | CENPL | 3.325 | 101 | NSL1 | 1.746 | 151 | PSMA4 | 1.128 |
| 2 | SPC25 | 28.289 | 52 | ORC1 | 3.265 | 102 | PSMC3 | 1.743 | 152 | PSMB3 | 1.114 |
| 3 | RPS27A | 27.516 | 53 | RFC2 | 3.244 | 103 | PSMA3 | 1.739 | 153 | CDKN1B | 1.108 |
| 4 | PSME1 | 21.896 | 54 | PPP2R5B | 3.204 | 104 | PMF1 | 1.733 | 154 | PPP2R5C | 1.108 |
| 5 | MCM10 | 15.916 | 55 | CDC20 | 3.180 | 105 | RPA2 | 1.720 | 155 | PPP2CA | 1.105 |
| 6 | CDT1 | 15.123 | 56 | MCM4 | 3.153 | 106 | PSMA5 | 1.707 | 156 | PPP2R1B | 1.103 |
| 7 | PSMC2 | 13.974 | 57 | RANBP2 | 3.138 | 107 | PSMD11 | 1.700 | 157 | CDK2 | 1.094 |
| 8 | CENPI | 13.831 | 58 | MCM6 | 3.083 | 108 | RPA1 | 1.698 | 158 | PPP1CC | 1.020 |
| 9 | AURKB | 12.728 | 59 | CENPN | 3.048 | 109 | CENPT | 1.646 | 159 | MAD1L1 | 0.989 |
| 10 | CDC6 | 12.029 | 60 | PPP2R5A | 2.991 | 110 | PSMB7 | 1.645 | 160 | PPP2R5D | 0.896 |
| 11 | BIRC5 | 9.943 | 61 | INCENP | 2.967 | 111 | PSMD12 | 1.644 | 161 | PSMD3 | 0.885 |
| 12 | GINS2 | 9.823 | 62 | MCM7 | 2.926 | 112 | PSMD1 | 1.634 | 162 | PSMB4 | 0.880 |
| 13 | SKA1 | 9.493 | 63 | POLD3 | 2.837 | 113 | PSMA2 | 1.616 | 163 | MAPRE1 | 0.868 |
| 14 | NDC80 | 9.282 | 64 | CASC5 | 2.796 | 114 | PAFAH1B1 | 1.608 | 164 | PSMB10 | 0.844 |
| 15 | CENPM | 8.938 | 65 | PSMA1 | 2.785 | 115 | PSMD6 | 1.595 | 165 | KIF2A | 0.822 |
| 16 | ERCC6L | 8.639 | 66 | KIF20A | 2.776 | 116 | POLD2 | 1.564 | 166 | KNTC1 | 0.814 |
| 17 | PSMD9 | 8.307 | 67 | RAD21 | 2.735 | 117 | PSMA7 | 1.533 | 167 | E2F3 | 0.773 |
| 18 | NUF2 | 7.717 | 68 | DNA2 | 2.724 | 118 | RFC4 | 1.528 | 168 | PSMD5 | 0.768 |
| 19 | CDC45 | 7.440 | 69 | LIG1 | 2.710 | 119 | POLA2 | 1.519 | 169 | STAG2 | 0.766 |
| 20 | KIF2C | 7.041 | 70 | CENPP | 2.706 | 120 | CLIP1 | 1.509 | 170 | POLD4 | 0.695 |
| 21 | NUDC | 7.037 | 71 | SKA2 | 2.696 | 121 | PSME2 | 1.509 | 171 | SEH1L | 0.626 |
| 22 | PRIM1 | 6.714 | 72 | SPC24 | 2.657 | 122 | POLE2 | 1.489 | 172 | PSMB2 | 0.624 |
| 23 | ZWINT | 6.504 | 73 | PPP2CB | 2.635 | 123 | PSMB1 | 1.467 | 173 | PSMC6 | 0.610 |
| 24 | CDCA8 | 6.473 | 74 | MAD2L1 | 2.634 | 124 | UBA52 | 1.466 | 174 | RFC5 | 0.548 |
| 25 | CENPK | 6.246 | 75 | FBXO5 | 2.612 | 125 | CCNA2 | 1.461 | 175 | PSME4 | 0.533 |
| 26 | MLF1IP | 5.954 | 76 | CDKN1A | 2.611 | 126 | PSMB5 | 1.451 | 176 | RCC2 | 0.502 |
| 27 | ZWILCH | 5.924 | 77 | RPA3 | 2.554 | 127 | ORC4 | 1.405 | 177 | CKAP5 | 0.496 |
| 28 | KIF18A | 5.346 | 78 | CCNA1 | 2.552 | 128 | PSMD14 | 1.402 | 178 | NDEL1 | 0.472 |
| 29 | GINS1 | 5.281 | 79 | CENPO | 2.513 | 129 | PSMD8 | 1.377 | 179 | CLASP1 | 0.438 |
| 30 | FEN1 | 5.251 | 80 | DSN1 | 2.460 | 130 | ORC3 | 1.366 | 180 | RANGAP1 | 0.436 |
| 31 | E2F1 | 4.845 | 81 | MCM3 | 2.456 | 131 | PSMB8 | 1.352 | 181 | CENPC1 | 0.417 |
| 32 | GORASP1 | 4.819 | 82 | NUP85 | 2.418 | 132 | PSMD13 | 1.344 | 182 | B9D2 | 0.320 |
| 33 | CENPA | 4.714 | 83 | PSMF1 | 2.379 | 133 | PSMD2 | 1.336 | 183 | PSMC1 | 0.121 |
| 34 | PLK1 | 4.472 | 84 | DBF4 | 2.332 | 134 | MIS12 | 1.320 | 184 | RPS27 | 0.020 |
| 35 | SGOL2 | 4.429 | 85 | MCM8 | 2.249 | 135 | XPO1 | 1.316 |  |  |  |
| 36 | BUB1 | 4.375 | 86 | POLE | 2.223 | 136 | PSMD4 | 1.282 |  |  |  |
| 37 | E2F2 | 4.169 | 87 | PSMC4 | 2.145 | 137 | GINS4 | 1.275 |  |  |  |
| 38 | ITGB3BP | 4.135 | 88 | SEC13 | 2.123 | 138 | NUP107 | 1.271 |  |  |  |
| 39 | TAOK1 | 3.893 | 89 | MCM2 | 2.109 | 139 | CENPH | 1.269 |  |  |  |
| 40 | PCNA | 3.872 | 90 | STAG1 | 1.914 | 140 | PSMA6 | 1.253 |  |  |  |
| 41 | ORC6 | 3.823 | 91 | PSMA8 | 1.895 | 141 | ORC5 | 1.211 |  |  |  |
| 42 | SMC3 | 3.778 | 92 | PPP2R1A | 1.836 | 142 | ZW10 | 1.187 |  |  |  |
| 43 | MCM5 | 3.717 | 93 | NUP133 | 1.833 | 143 | PRIM2 | 1.182 |  |  |  |
| 44 | CENPQ | 3.709 | 94 | SMC1A | 1.804 | 144 | PSMB9 | 1.167 |  |  |  |
| 45 | SGOL1 | 3.628 | 95 | PSMC5 | 1.799 | 145 | PSMB6 | 1.157 |  |  |  |
| 46 | RFC3 | 3.606 | 96 | PPP2R5E | 1.790 | 146 | NUP43 | 1.156 |  |  |  |
| 47 | CDC7 | 3.542 | 97 | ORC2 | 1.776 | 147 | POLA1 | 1.147 |  |  |  |
| 48 | KIF23 | 3.485 | 98 | PSMD10 | 1.773 | 148 | PSMD7 | 1.138 |  |  |  |
| 49 | APITD1 | 3.353 | 99 | NUP37 | 1.771 | 149 | RB1 | 1.131 |  |  |  |
| 50 | BUB1B | 3.351 | 100 | BUB3 | 1.767 | 150 | AHCTF1 | 1.129 |  |  |  |

